# FHL5 controls vascular disease-associated gene programs in smooth muscle cells

**DOI:** 10.1101/2022.07.23.501247

**Authors:** Doris Wong, Gaëlle Auguste, Christian L. Lino Cardenas, Adam W. Turner, Yixuan Chen, Lijiang Ma, R. Noah Perry, Redouane Aherrahrou, Maniselvan Kuppusamy, Chaojie Yang, Jose Verdezoto Mosquera, Collin J. Dube, Mohammad Daud Khan, Meredith Palmore, Maryam Kavousi, Patricia A. Peyser, Ljubica Matic, Ulf Hedin, Ani Manichaikul, Swapnil K. Sonkusare, Mete Civelek, Jason C. Kovacic, Johan L.M. Björkegren, Rajeev Malhotra, Clint L. Miller

## Abstract

**Background:** Genome-wide association studies (GWAS) have identified hundreds of loci associated with common vascular diseases such as coronary artery disease (CAD), myocardial infarction (MI), and hypertension. However, the lack of mechanistic insights for a majority of these loci limits translation of these findings into the clinic. Among these loci with unknown functions is *UFL1-FHL5* (chr6q16.1), a locus that reached genome-wide significance in a recent CAD/MI GWAS meta-analysis. In addition to CAD/MI, *UFL1-FHL5* is also implicated to coronary calcium, intracranial aneurysm, and migraine risk, consistent with the widespread pleiotropy observed among other GWAS loci.

**Methods:** We apply a multimodal approach leveraging statistical fine-mapping, epigenomic profiling, and imaging of human coronary artery tissues to implicate Four-and-a-half LIM domain 5 (*FHL5)* as the top candidate causal gene. We unravel the molecular mechanisms of the cross-phenotype genetic associations through *in vitro* functional analyses and epigenomic profiling experiments.

**Results:** We prioritized FHL5 as the top candidate causal gene at the *UFL1-FHL5* locus through eQTL colocalization methods. FHL5 gene expression was enriched in the SMC and pericyte population in human artery tissues with coexpression network analyses supporting a functional role in regulating SMC contraction. Unexpectedly, under procalcifying conditions, FHL5 overexpression promoted vascular calcification and dysregulated processes related to extracellular matrix organization and calcium handling. Lastly, by mapping FHL5 binding sites and inferring FHL5 target gene function using artery tissue gene regulatory network analyses, we highlight regulatory interactions between FHL5 and downstream CAD/MI loci, such as *FOXL1* and *FN1* that have roles in vascular remodeling.

**Conclusion:** Taken together, these studies provide mechanistic insights into the pleiotropic genetic associations of *UFL1-FHL5*. We show that FHL5 mediates vascular disease risk through transcriptional regulation of downstream vascular remodeling loci. These *trans*-acting mechanisms may account for a portion of the heritable risk for complex vascular diseases.

## Introduction

Vascular diseases, such as atherosclerosis and aneurysm, encompass a wide range of disorders that affect blood vessels and perturb blood flow to critical tissues. Despite the heritability of these common pathologies, many of the associated genetic risk factors are unknown. To address this clinical gap, genome-wide association studies (GWAS) have been used to uncover the genetic basis for many complex traits. In the case of coronary artery disease (CAD), which is among the leading causes of death worldwide, GWAS have identified over 200 loci^1^. Interestingly, a majority of these loci harbor genes that function independently of lipid metabolism and other classic risk factors. These studies highlight additional pathways, such as vascular remodeling and inflammation that directly contribute to CAD pathogenesis and prioritize novel therapeutic targets^2,3^.

As demonstrated by lineage tracing and single-cell RNA sequencing (RNA-seq) studies in mice and humans, smooth muscle cells (SMCs) play a key role in both the development and progression of atherosclerosis^4,5^. SMCs undergo phenotypic transitions to give rise to diverse cell populations that drive atherosclerosis progression in early stages and contribute to fibrous cap stability in advanced lesions^6^. Intimal SMCs that acquire mesenchymal and ultimately osteoblast-like phenotypes deposit calcium-mineral in the collagenous matrix^7^. These remodeling events coupled with arterial calcification serve as strong predictors of adverse cardiovascular events^8^. Untangling the complex contribution of SMCs to CAD using human genetics may inform novel drug candidates targeting these primary disease processes in the vessel wall.

The most recent CAD and myocardial infarction (MI) meta-analysis of the combined UK Biobank and CARDIoGRAMplusC4D cohorts identified a genome-wide significant association of the *UFL1-FHL5* locus (chr6q16.1)^9^. In addition to CAD/MI, *UFL1-FHL5* is also associated with multiple vascular pathologies, including hypertension^10^, intracranial aneurysm^11^, and migraines^12^, similar to another pleiotropic locus *PHACTR1-EDN1*^*13*^. The lead CAD/MI single nucleotide polymorphism (SNP), rs9486719 resides in the first intron of *FHL5*. FHL5 is a member of the four-and-a-half LIM (FHL) domain family of cofactors, which also includes structurally similar proteins, FHL1, FHL2, and FHL3^14^. Among the FHL family, FHL5 is the most understudied, which may be attributed to its low expression *in vitro* and limited availability of suitable animal models. Early functional studies have focused on its expression in germ cells^15,16^, however FHL5 was recently linked to intimal hyperplasia in aortic SMCs by activating CREB target genes^17,18^.

Here, we present a comprehensive study that investigates the molecular underpinnings of the genetic association of the *UFL1-FHL5* locus with CAD/MI and other vascular pathologies. We identify an FHL5-regulated transcriptional network that contributes to the maladaptive extracellular matrix (ECM) remodeling and vascular tone defects associated with clinical disease risk.

## Methods

A detailed and expanded methods section can be found in the Supplemental Material. A list of key reagents can be found in the Major Resources Table located in the Supplementary Materials section.

### Statistical analysis

Data in bar graphs are presented with mean+/- standard error of mean (SEM) with each point represented as an individual replicate. Data in box plots are presented with lines denoting the 25th, median and 75th percentile with each point representing an individual donor. Pairwise comparisons were made using the student’s t test or Wilcoxon rank test as appropriate. Comparisons between more than two groups were assessed using a one-way ANOVA test or Kruskal-Wallis test. The normality of the data was assessed using the Shapiro-Wilkes test, with P > 0.05, supporting a normal distribution. For each of these analyses, we considered P < 0.05 as significant.

To identify differentially expressed genes, we used the false discovery rate (FDR) adjusted P< 0.05 threshold and log2FoldChange > 0.6. Heatmaps were created using the pheatmap package and represent normalized expression (Z-score) for genes, scaled across each row. Gene ontology enrichment analyses were performed relative to all expressed genes using Fisher’s Exact Test, with a significant threshold of 5% FDR.

### Data Availability

All raw and processed CUT&RUN and RNA-seq datasets are made available on the Gene Expression Omnibus (GEO) database (accession: GSE201572). All other datasets are publicly available and detailed in the Major Resources Table Section or Supplemental Materials section. All custom scripts used are available at https://github.com/MillerLab-CPHG/FHL5_Manuscript. All software tools used in this study are publicly available and full names and versions are provided in Supplementary Materials.

### Ethics statement

All research described herein complies with ethical guidelines for human subjects research under approved Institutional Review Board (IRB) protocols at Stanford University (#4237) and the University of Virginia (#20008), for the procurement and use of human tissues and information, respectively.

## Results

### FHL5 is the top candidate causal gene associated with CAD/MI risk

The lead variant, rs9486719, tagging the *UFL1-FHL5* locus (chr6q16.1) is associated with CAD (*P* = 1.1E-8) and MI (*P* = 6.8E-10) risk (**Figure 1A** and **Figure S1A**), as reported in the combined genome-wide association study (GWAS) meta-analysis of CARDIoGRAMplusC4D and UK Biobank (UKBB) data^1,9^. The lead MI SNP, rs9486719 is also associated with intermediate traits predictive of cardiovascular adverse events, such as blood pressure^19,20^ and hypertension^10^ **(Figure 1B)**. We did not observe genetic associations of rs9486719 with lipid metabolism or other traditional CAD risk factors (e.g. type 2 diabetes or obesity) in the PhenoScanner database^21,22^ (**Figure 1C** and **Table S1)**, implicating heritable CAD risk at this locus acting primarily in the vessel wall.

**Figure 1.**
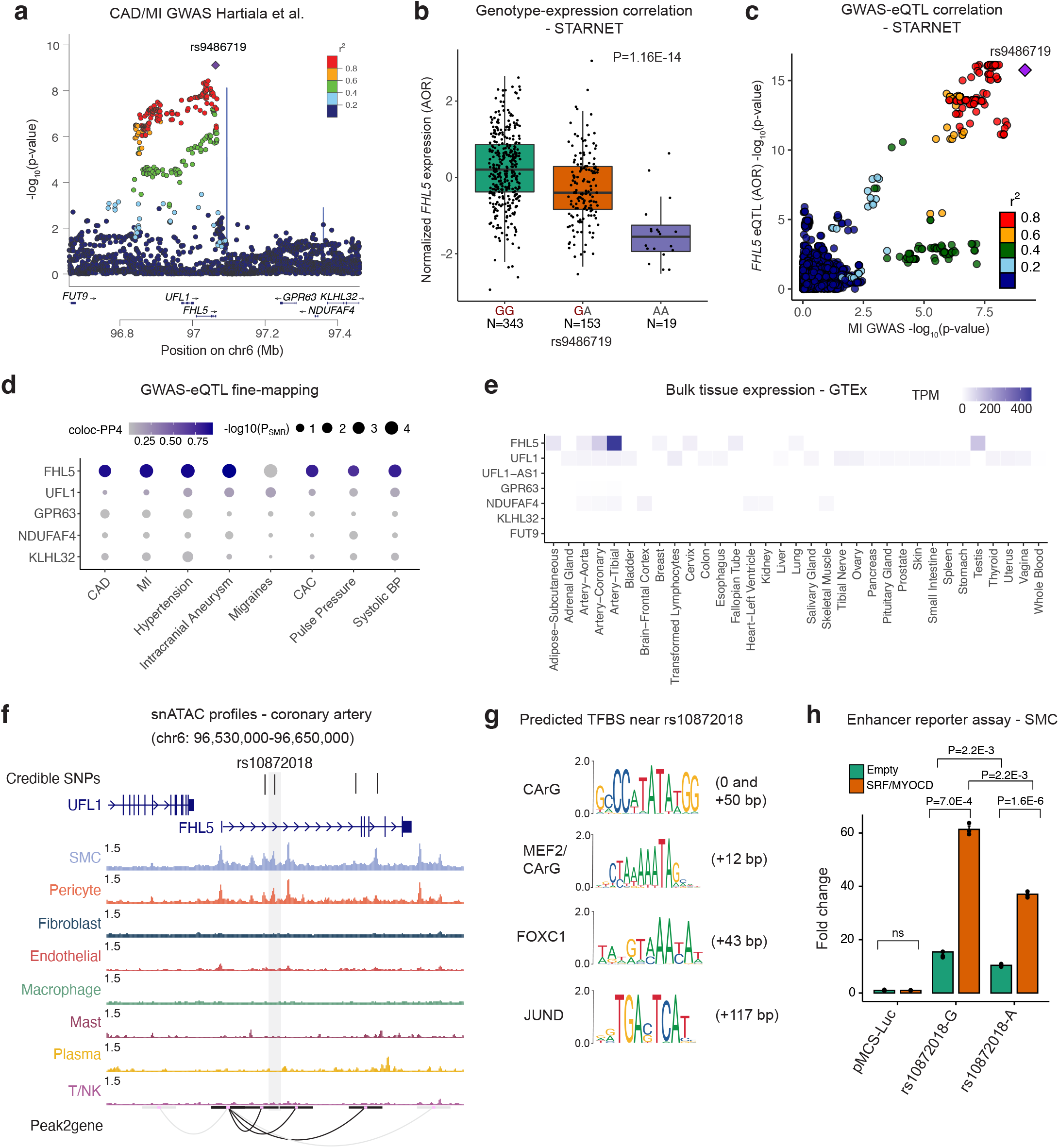
*FHL5* is the top candidate causal gene at the *UFL1*-*FHL5* locus associated with increased CAD/MI risk. **(a)** LocusZoom plot showing the association of *UFL1-FHL5* locus with myocardial infarction (MI) in the combined UKBB and CARDIoGRAMplusC4D meta-analysis. **(b)** Association of rs9486719 with *FHL5* gene expression in STARNET aortic tissue with the CAD risk allele (rs9486719-G) associated with increased FHL5 gene expression. Box plots represent the median with the box spanning the first and third quartiles and whiskers as 1.5 times IQR. **(c)** Locuscompare plot showing the associations of top *FHL5* STARNET aortic tissue *cis*-eQTLs with MI. **(d)** Summary of SMR and coloc analyses nominating *FHL5* as top candidate causal gene for vascular diseases and cardiovascular risk factors. The size of the dot reflects the -log10(pSMR) and the intensity of purple color represents the posterior probability of colocalization. **(e)** Normalized gene expression as transcripts per million (TPM) for all genes at the *UFL1-FHL5* locus across GTEx tissues. **(f)** Genome browser view of human coronary artery snATAC-seq peaks showing overlap between CAD 95% credible set of SNPs (highlighting top candidate rs10872018) with putative enhancers correlated with the *FHL5* promoter through peak2Gene analyses. **(g)** Predicted transcription factor binding sites (TFBS) at or around rs10872018 determined from JASPAR 2022. Distance (bp) of motif sequence is also shown relative to the rs10872018 SNP. **(h)** Luciferase reporter assay in A7r5 SMCs comparing rs10872018 allele-specific enhancer activity co-transfected with empty vector (control) or *SRF* and *MYOCD* expression constructs.

rs9486719 is located in the first intron of *FHL5*, and similar to the majority of GWAS variants likely modulates gene expression to influence disease risk^23,24^ (**Figure 1A)**. To prioritize the most likely target gene(s) at this locus, which includes 7 genes, we leveraged *cis*-expression quantitative trait loci (*cis*-eQTLs) in cardiometabolic tissues from Stockholm-Tartu Atherosclerosis Reverse Network Engineering Task (STARNET)^25^. This SNP was strongly associated with *FHL5* gene expression in both mammary arteries (MAM) with subclinical atherosclerotic disease, and aortic root tissues (AOR) with atherosclerosis (**Figure 1B**). These results were consistent with GTEx eQTLs, in which rs9486719 was significantly associated with *FHL5* expression in aorta (P=2.3E-7) (**Figure S2A**). Importantly, *FHL5* eQTLs colocalized with CAD SNPs in aortic tissues, compared to eQTLs for neighboring genes (e.g., *UFL1*) (**Figure 1C**). In order to systematically prioritize the GWAS effector genes at the *UFL1-FHL5* locus, we employed two complementary and independent fine-mapping methods, *coloc*^*26*,27^ *and* Summary-level based Mendelian Randomization (SMR)^28,29^. By integrating GTEx and STARNET artery tissue eQTLs for all the genes at the *UFL1-FHL5* locus with GWAS summary statistics, we consistently identified *FHL5* as the top candidate causal gene for CAD/MI and other common vascular traits (**Figure 1D** and **Table S2**). This was consistent with the tissue distribution of *FHL5* gene expression, which was highly enriched in artery tissues in GTEx (**Figure 1E**) and STARNET (**Figure S2B**) compared to other genes at the locus. Together, these analyses suggest that the *FHL5* risk allele (rs9486719-G) increases CAD/MI risk by increasing *FHL5* gene expression in artery tissue (**Figure S2C)**.

### Epigenomic based fine-mapping of UFL1-FHL5 locus in human artery tissue

Next, we leveraged human coronary artery epigenomic profiles to resolve candidate causal variants at the *UFL1-FHL5* locus. The MI 95% credible set determined from Probabilistic Annotation Integrator v3 (PAINTOR)^30^ using human coronary artery SMC and pericyte open chromatin profiles (snATAC-seq) as prior functional annotation weights consisted of 4 SNPs **(Figure S3A** and **Table S3**). By overlapping variants in this credible set with SMC Peak2Gene linkages^31^, we prioritized rs10872018, a SNP located in an active *FHL5* regulatory element *in vivo*. (**Figure 1F**). These interactions were supported by the Activity-by-Contact Model (ABC)^32^ generated using ENCODE human coronary artery tissue ATAC-seq, H3K27ac, and HiChIP data, which further prioritized rs10872018 as the most likely causal variant (**Table S4**). This variant was identified as one of the top eQTLs in STARNET human vascular tissues (**Figure S3B**).

Since trait-associated SNPs are predicted to alter regulatory elements^33^, we scanned the genomic sequence +/- 100bp of rs10872018 for putative transcription factor binding sites and identified multiple conserved CArG boxes^34^ (**Figure S3C**). The alternate allele (rs10872018-A) disrupts a non-canonical CArG box motif, which likely perturbs binding of the SRF-Myocardin (MYOCD) transcriptional complex, a well-characterized regulator of SMC differentiation^35,36^. We validated this putative upstream regulatory mechanism using allele-specific enhancer luciferase reporter assays in A7r5 aortic SMCs. The risk allele (rs10872018-G) increased luciferase activity relative to the non-risk allele (rs10872018-A) consistent with the *FHL5* eQTL direction in human artery tissues (**Figure 1G** and **Figure S3B)**. SRF and MYOCD overexpression further potentiated the luciferase activity in an allele-specific manner (**Figure 1G**). Together, these results demonstrate that altered SRF-MYOCD binding due to rs10872018 may partially explain the observed eQTL effects in human artery tissues during disease.

### FHL5 is highly enriched in contractile mural cells in coronary arteries

To confirm expression of FHL5 protein in human artery tissues and identify its endogenous localization, we performed immunofluorescence on sections of human coronary arteries with subclinical atherosclerosis. FHL5 protein was enriched in the SMC-containing medial layer as well as the intima, colocalizing with F-actin positive cells. (**Figure 2A**). Consistent with previous reports as a transcriptional regulator^37^, we observed nuclear localization of FHL5 in both the medial and intimal layers, but also perinuclear/cytoplasmic localization. To identify the cell-type specific expression profile, we queried a human coronary artery scRNA-seq dataset from 4 donors^6,38^, which revealed *FHL5* gene expression enriched in the mural cells (SMCs and pericytes) (**Figure 2B)**. In fact, *FHL5* was one of the most specific markers identified in the SMC cluster along with well-established regulators of SMC contraction, e.g. *LMOD1* and *MYOCD* (**Figure S4A**). Similar results were observed in an advanced carotid artery single cell RNA-seq dataset^39^ (**Figure S4B**). Lastly, we corroborated these results in a single-nucleus ATAC-seq (snATAC-seq) dataset of 41 human coronary arteries^40^, in which *FHL5* had the highest gene score in the SMC and pericyte clusters, similar to *LMOD1* (**Figure 2C** and **Figure S4C**).

**Figure 2.**
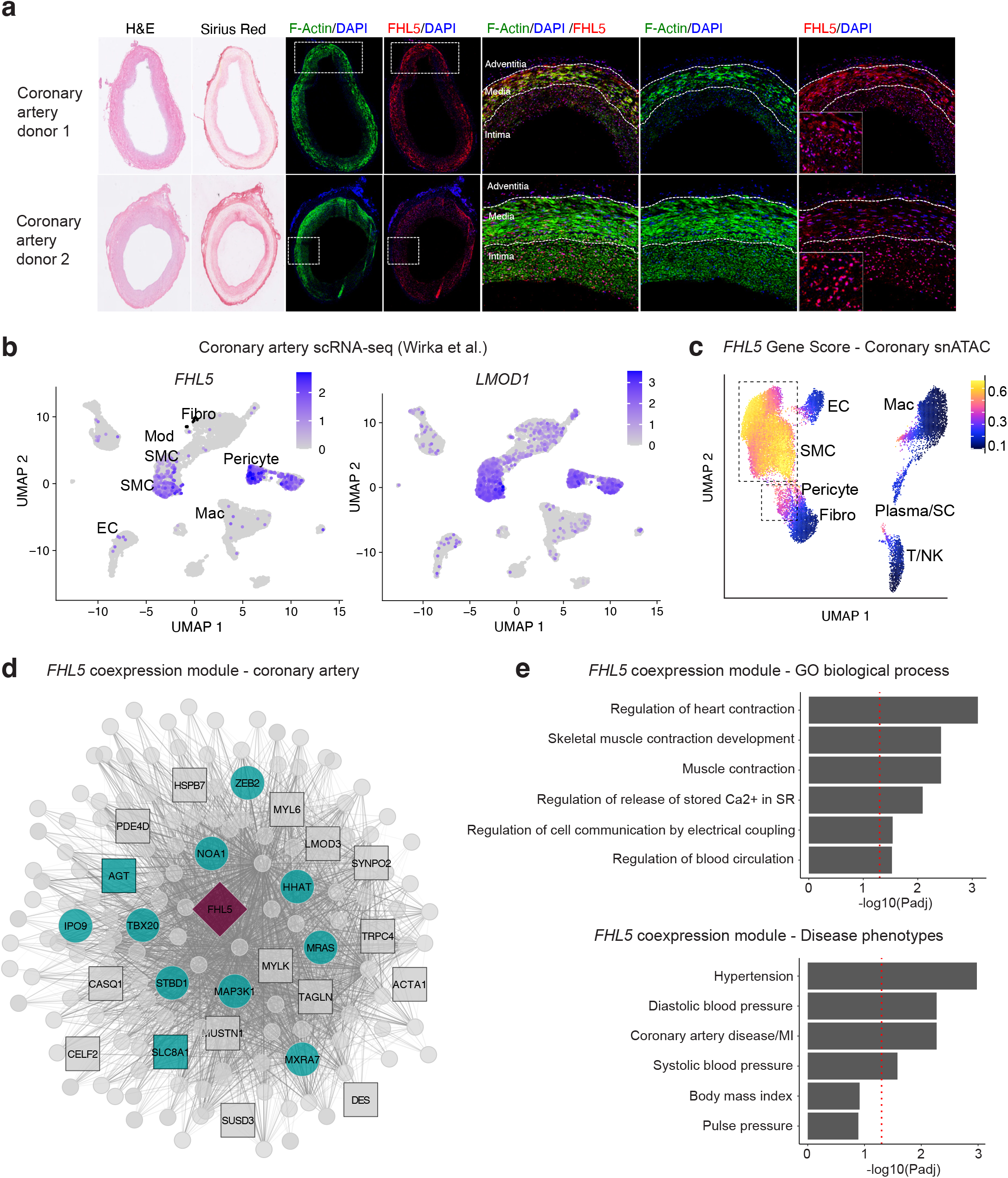
FHL5 expression is enriched in SMCs and pericytes in human coronary arteries. **(a)** Representative Immunofluorescence (IF) of FHL5 protein (red) colocalizing with F-actin (green) immunopositive SMCs in the medial and intimal layers of human subclinical atherosclerotic coronary arteries (n=8 unique donors). **(b)** UMAP visualization of *FHL5* and LMOD1 gene expression in different human coronary artery cell types from Wirka et al ^6^. **(c)** UMAP visualization of human coronary artery snATAC-seq cell clusters colored according to *FHL5* gene score calculated in ArchR. **(d)** Undirected *FHL5* coexpression network identified in human coronary arteries using weighted gene co-expression network analysis (WGCNA). Each node represents a gene and the lines connecting 2 nodes are weighted according to degree of correlation. The teal color highlights module genes associated with CAD/MI or blood pressure. The square nodes represent module genes that regulate SMC contraction. *FHL5* is placed at the center and depicted as a red diamond. **(e)** Enrichment of gene ontology biological processes (GO-BP) and **(f)** cardiometabolic disease phenotypes in the *FHL5* module protein-coding genes. The highlighted terms are a subset of the full ranked list found in Supp Table 6.

To gain insights into the functional role of FHL5 using a systems biology approach, we performed iterative Weighted Gene Co-expression Network Analysis (WGCNA)^41,42^ on transcriptomic data from 148 human coronary artery donors. After removing outliers and lowly expressed genes, we identified *FHL5* in the light-green module, which included key SMC contractile mediators, such as *MYLK, ACTA2, and TAGLN* **(Figure 2D** and **Table S5A)**. This module was enriched in SMC processes, such as muscle contraction (GO:0006936) and regulation of cell communication by electrical coupling (GO:0010649) as well as cardiometabolic GWAS candidate genes annotated in the Cardiovascular Disease Knowledge Portal (**Figure 2E** and **Table S6**). In further support of this link to vascular disease risk through regulation of SMC functions, FHL5 was identified as a key driver in module 152, a cross-tissue gene regulatory network enriched in ECM and CAD candidate genes from the STAGE cohort^43^. The distinct enrichment in pathways among these two vascular tissues may reflect differences in the atherosletic lesion stage.

To further characterize *FHL5* regulatory interactions *in vivo*, we constructed a Bayesian GRN incorporating STARNET aortic tissue eQTL data as priors (**Figure S5A**). A key driver analysis (KDA) of this network supported the regulatory potential of *FHL5* which was predicted to function upstream of 2 pulse pressure genes, *ACTG* and *MUSTN1*^*19*^ that have established roles in SMC contraction (**Figure S5B** and **Table S5B**). Notably, given its link to MI in a Japanese population^44^, *ITIH3* was the key driver gene in this subnetwork. These integrative analyses suggest that this *FHL5* GRN related to SMC contractility may contribute to pleiotropic association of FHL5 with common vascular diseases.

### FHL5 expression increases SMC contraction and calcification

To overcome the limitations in replicative senescence in primary HCASMCs, we generated an immortalized coronary artery SMC line via overexpression of hTERT^45^. This immortalized cell line (HCASMC-hTERT) maintained protein expression of differentiated SMC markers and closely resembled primary HCASMCs at the transcriptomic level. (**Figure S6A** through **S6C**). Despite robust expression in intact arteries, *FHL5* is potently downregulated *in vitro*, similar to the downregulation of other SMC contractile markers (**Figure S7A)**. Therefore, to investigate its role in SMCs, we overexpressed either wildtype FHL5 or FHL5-NLS (FHL5 with a C-terminal nuclear localization signal) and confirmed protein expression (**Figure 3A**). The expression level of *FHL5* was physiological and comparable to endogenous levels in human coronary arteries (**Figure 3B)**. Motivated by our coexpression network analyses, we functionally validated the role of FHL5 in regulating SMC contractile pathways. Since calcium is a critical initiator of contraction, we also quantitated intracellular calcium levels. We observed increased SMC contraction and elevated calcium levels in the FHL5-NLS cells relative to HA control cells. While not statistically significant, FHL5 overexpressing cells trended in the same direction (**Figure 3C** and **3D**).

**Figure 3.**
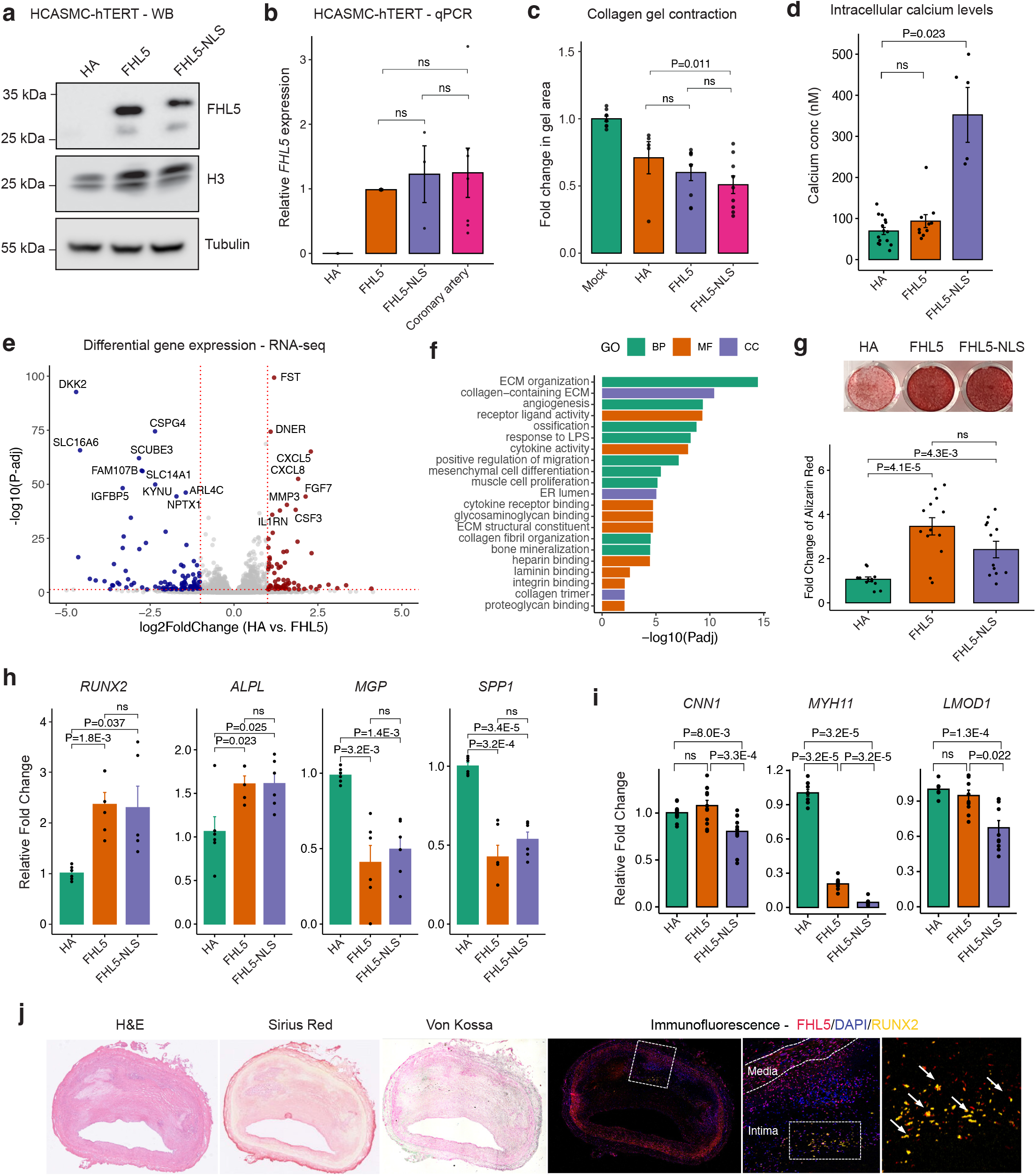
*FHL5* regulates SMC contraction and calcification. **(a)** Western blot confirming overexpression of FHL5 and FHL5-NLS in HCASMC-hTERT cell lines. **(b)** Relative expression of FHL5 in HCASMC-hTERT (HA, FHL5, FHL5-NLS) compared to endogenous levels of FHL5 in human coronary arteries (n=3 donors) determined using qPCR. Expression was normalized to levels of *GAPDH*. **(c)** Relative change in collagen gel area (mm^2^) in SMC contraction assay under basal conditions (n=4 independent biological replicates). **(d)** Quantification of intracellular calcium concentrations following stimulation with 10µm phenylephrine (n=4 biological replicates). **(e)** Volcano plot showing differentially expressed genes (DEGs) following FHL5 overexpression in HCASMC-hTERT. Red points represent upregulated genes and blue points represent downregulated genes. For clarity, genes with log2FoldChange > 5 and log2Foldchange <-5 are not represented. The full DEG list is provided in Supplementary Table 8. **(f)** GO enrichment analysis of FHL5 DEGs showing top over-represented terms for biological process (BP), molecular function (MF), and cell compartment (CC). These highlighted terms were selected from the full ranked list found in Supplementary Table 9. **(g)** Quantification (OD405) and representative image of alizarin red staining 21 days post treatment of HCASMC-hTERT in osteogenic media. **(h)** Relative expression of vascular calcification activators (*RUNX2, ALPL)* and inhibitors (*MGP, SPP1)* 14 days post treatment in osteogenic media. **(i)** Relative expression of SMC markers 14 days post treatment in osteogenic media. **(j)** Representative immunofluorescence staining of human coronary arteries showing FHL5 (red) colocalization with RUNX2(yellow) in the intima layer near regions of calcium deposition (n=4 independent donors). Adjacent sections subjected to histology staining for hematoxylin & eosin (H&E), collagen deposition (Sirius Red), and calcification (Von Kossa). All error bars represent mean+/-SEM. P-values determined from paired Student’s t-test. Individual points reflect replicates from at least n=3 independent experiments. ns: non-significant.

Next, to investigate the molecular mechanisms contributing to FHL5 regulation of SMC phenotypes, we performed bulk RNA-seq on HCASMC-hTERT expressing HA, FHL5 or FHL5-NLS constructs. Top differentially expressed genes had mainly concordant direction of effects between FHL5 and FHL5-NLS samples (**Figure S7B** through **S7D** and **Table S7**). When comparing FHL5 versus HA cells, we identified 377 differentially expressed genes (log2FoldChange > 0.6 and FDR < 0.05), of which 191 and 186 genes were upregulated and downregulated respectively (**Figure 3E** and **Table S7A)**. Top upregulated genes included various metalloproteinases (*MMP1, MMP3, MMP10*) and vessel wall matrisome proteins (*DCN, COL5A3, ANGPTL4*). We noted that FHL5-mediated differentially expressed genes (DEGs) were overrepresented in vascular remodeling pathways, such as cytokine activity/inflammation, ECM organization and ossification (**Figure 3F** and **Table S8**). Interestingly, FHL5 DEGs were also enriched with vascular and inflammatory disease candidate genes (**Figure S7E**).

Based on the perturbation of vascular remodeling and ossification pathways supported by RNA-seq, we next hypothesized that FHL5 may regulate vascular calcification to mediate CAD/MI risk. To this end, we treated SMCs with an osteogenic cocktail as done previously^46,47^. We observed increased mineral deposition quantified by alizarin red staining upon FHL5 and FHL5-NLS overexpression (**Figure 3G**). This increased calcification correlated with increased expression of osteogenic activators, *RUNX2* and *ALPL* and reduced expression of osteogenic inhibitors, *MGP* and *SPP1* (**Figure 3H)**. Consistent with promoting SMC phenotypic transitions toward an osteogenic state, FHL5 and FHL5-NLS overexpression also coincided with downregulated expression of SMC markers, *LMOD1, MYH11*, and *CNN1* (**Figure 3I)**. Lastly, to validate the role of FHL5 in increasing vascular calcification, we immunostained sections of human coronary artery plaques. In accordance with our *in vitro* findings, we observed colocalization between FHL5 and RUNX2, a transcriptional activator of osteogenic differentiation in close proximity to intimal calcium deposits (**Figure 3J**). These results were consistent with the direction of effect of the *FHL5* genetic association with coronary artery calcification (CAC), where the CAC risk allele was associated with increased *FHL5* gene expression. Together, these analyses suggest that FHL5 promotes a shift towards the SMC osteogenic phenotypic state in atherosclerotic lesions to increase CAD/MI risk.

### FHL5 serves as a SMC cofactor to regulate disease-associated ECM interactions

In order to further decipher the direct transcriptional network regulated by FHL5 in SMCs under basal conditions, we used the Cleavage Under Targets and Release Using Nuclease (CUT&RUN) method to map genome-wide binding sites^48,49^. In total, we identified 17,201 FHL5 binding sites (qvalue < 0.01) that map to 6,776 unique genes, with a majority of binding sites located in annotated promoters and intronic regions (**Figure 4A** and **Figure S8A**)^50,51^. The active enhancer H3K27ac and promoter mark, H3K4me3 signals were enriched around the center of FHL5 peaks (**Figure 4B, 4C** and **Figure S8B through S8D**), with about 75% of binding sites overlapping primary HCASMC ATAC-seq peaks^52^ (**Figure S8E**).

**Figure 4.**
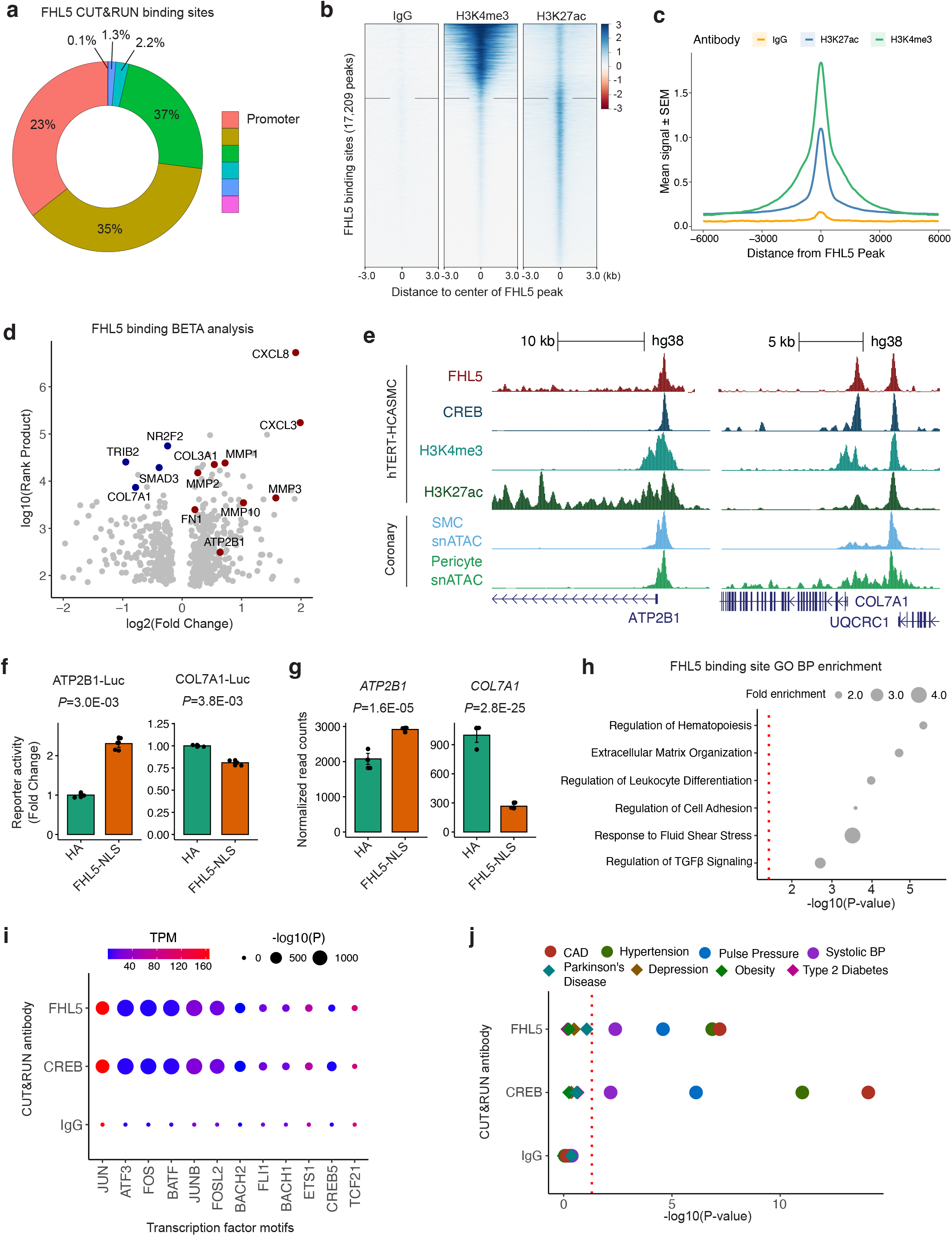
FHL5 serves as a cofactor for the transcription factor, CREB, to regulate ECM organization in SMCs. **(a)** Overlap of FHL5 peaks identified from CUT&RUN with genomic features. **(b)** Heatmap showing distribution of active chromatin histone marks (H3K4me3 and H3K27ac) +/-3kb from the center of FHL5 peaks, compared to IgG control. **(c)** Density plot showing genome-wide enrichment of active chromatin histone marks +/- 6kb from the center of FHL5 peaks. **(d)** Results of Binding and Expression Target Analysis (BETA) using FHL5 CUT&RUN peaks and FHL5 differentially expressed genes (DEGs) as input. Top candidate direct target genes highlighted in red, or blue based on upregulated and downregulated expression, respectively. Log normalized rank sum product score reflects the likelihood of direct transcriptional regulation for each gene. **(e)** UCSC genome browser tracks showing FHL5 binding sites at the *ATP2B1* and *COL7A1* loci, highlighting candidate regulatory elements overlapping CREB binding sites and H3K27ac and H3K4me3 peaks. These regions also overlapped accessible chromatin SMCs and pericytes determined from human coronary artery snATAC-seq. **(f)** Relative change in luciferase reporter activity of *ATP2B1* and *COL7A1* enhancer regions in FHL5 overexpressing SMCs. **(g)** Relative change in RNA expression of *ATP2B1* and *COL7A1* in SMCs overexpressing FHL5, shown as normalized read counts. **(h)** Top GO-BP overrepresented in FHL5 target genes identified from GREAT analysis of FHL5 peaks. These highlighted terms were selected from the full ranked list in Supp Table 10. **(i)** Top transcription factor motifs enriched in FHL5, CREB, and IgG peaks identified from HOMER known motif analysis (dot size) as well as normalized (transcripts per million: TPM) expression level of the corresponding transcription factors in SMCs (color scale). **(j)** GREGOR analysis showing enrichment of vascular trait GWAS risk variants in FHL5, CREB, and IgG binding sites.

To examine the functional consequences of FHL5 binding on transcription, we integrated genome-wide FHL5 binding sites with differential expression analysis using the Binding and Expression Target Analysis (BETA) tool^53^. We observed substantial overlap between FHL5 DEGs and FHL5 peaks (P=6.3E-42, Fisher’s Exact Test), with a majority (51%) of these differentially expressed genes harboring at least 1 FHL5 binding site near its transcription start site. FHL5 directly upregulated key genes involved in vascular remodeling, including contractile genes, metalloproteinases, and ECM components (**Figure 4D** and **Table S9**). For example, we identified FHL5 binding at the 5’ regulatory elements near the *ATP2B1*, a CAD associated gene involved in calcium ion homeostasis, and at the gene promoter of *COL7A1*, a component of the basement membrane (**Figure 4E**). We validated the activity of these regulatory elements at the *ATP2B1* and *COL7A1* promoter in SMCs. FHL5 overexpression increased luciferase activity and correlated with increased *ATP2B1* expression (**Figure 4F**). In contrast, FHL5 overexpression reduced the luciferase activity of the *COL7A1* enhancer and *COL7A1* gene expression, supporting the gene-specific activating or repressive role of FHL5 (**Figure 4G**). Gene ontology enrichment analysis of these high confidence target genes revealed FHL5 mediated direct regulation of pathways related to ECM organization and cell adhesion. We complemented this approach with Genomic Regions ENrichment of Annotations Tool (GREAT)^54^ analysis **(Figure 4H** and **Table S10)**. FHL5 binding sites were proximal to genes related to inflammation, ECM organization and cell adhesion. These enrichments mirror the BETA results and further highlight the regulatory interactions between FHL5 and downstream vascular remodeling genes in mural cells.

Since FHL5 has not been reported to bind DNA directly, we performed motif enrichment analysis on genome-wide FHL5 peaks to identify candidate transcription factor binding partners, which revealed enrichment of AP1 and cAMP-response element (CRE) motifs within these peaks (**Figure 4I**). Regulatory elements proximal to upregulated genes after FHL5 overexpresison were also enriched in AP1 motifs. Interestingly, consistent with previous studies, 48% of FHL5 binding sites overlapped CREB binding sites, which was also strikingly enriched in AP1 binding motifs as well (**Figure S8E** and **S8G**). We further explored additional potential transcription factor binding partners using the Locus Overlap Analysis (LOLA) enrichment tool^55^ and assessed the enrichment of FHL5 binding sites in the ENCODE collection of transcription factor ChIP-seq datasets. Relative to the HCASMC ATAC-seq peak set, we observed significant enrichment of Pol2, AP1 members, (c-Fos and c-Jun), CREB and its common cofactor, p300 (**Figure S8F**). We validated the role of FHL5 as a cofactor for CREB through a cAMP response element (CRE) luciferase reporter assay. Relative to the HA control, FHL5 overexpression upregulated CRE activity (**Figure S8H**).

### FHL5 regulates a transcriptional network that contributes to CAD/MI risk by modulating SMC functions and vascular remodeling processes

As a transcriptional regulator of CAD-relevant pathways in the vessel wall, we next hypothesized that FHL5 regulation of downstream target genes contribute to the mechanistic link between FHL5 and vascular disease risk. To this end, we evaluated the enrichment of vascular disease GWAS SNPs in FHL5 binding sites using the Genomic Regulatory Elements and Gwas Overlap algoRithm (GREGOR) software package. Relative to a matched set of random SNPs^56,57^. FHL5 and CREB binding sites were highly enriched for CAD, MI, and blood pressure risk variants (**Figure 4J**), compared to non-vascular traits, further emphasizing the specific contribution of FHL5 to the heritability of common vascular diseases.

To further support this hypothesis and extend our findings to human artery tissues, we assessed the distal effects of *FHL5* gene expression in STARNET cardiometabolic tissues. The top *FHL5 cis*-eQTL rs10872018 was associated with the expression of 743 and 568 eGenes in aortic and mammary artery tissues respectively at nominal significance (P < 0.05) (**Table S11**). These eGenes included key matrisome genes, such as *FN1, COL4A4*, and *LUM*. Integration of *trans*-eQTL target genes from STARNET artery tissues and FHL5 binding sites identified a network of downstream genes that harbor CAD risk variants (**Figure 5A)**.

**Figure 5.**
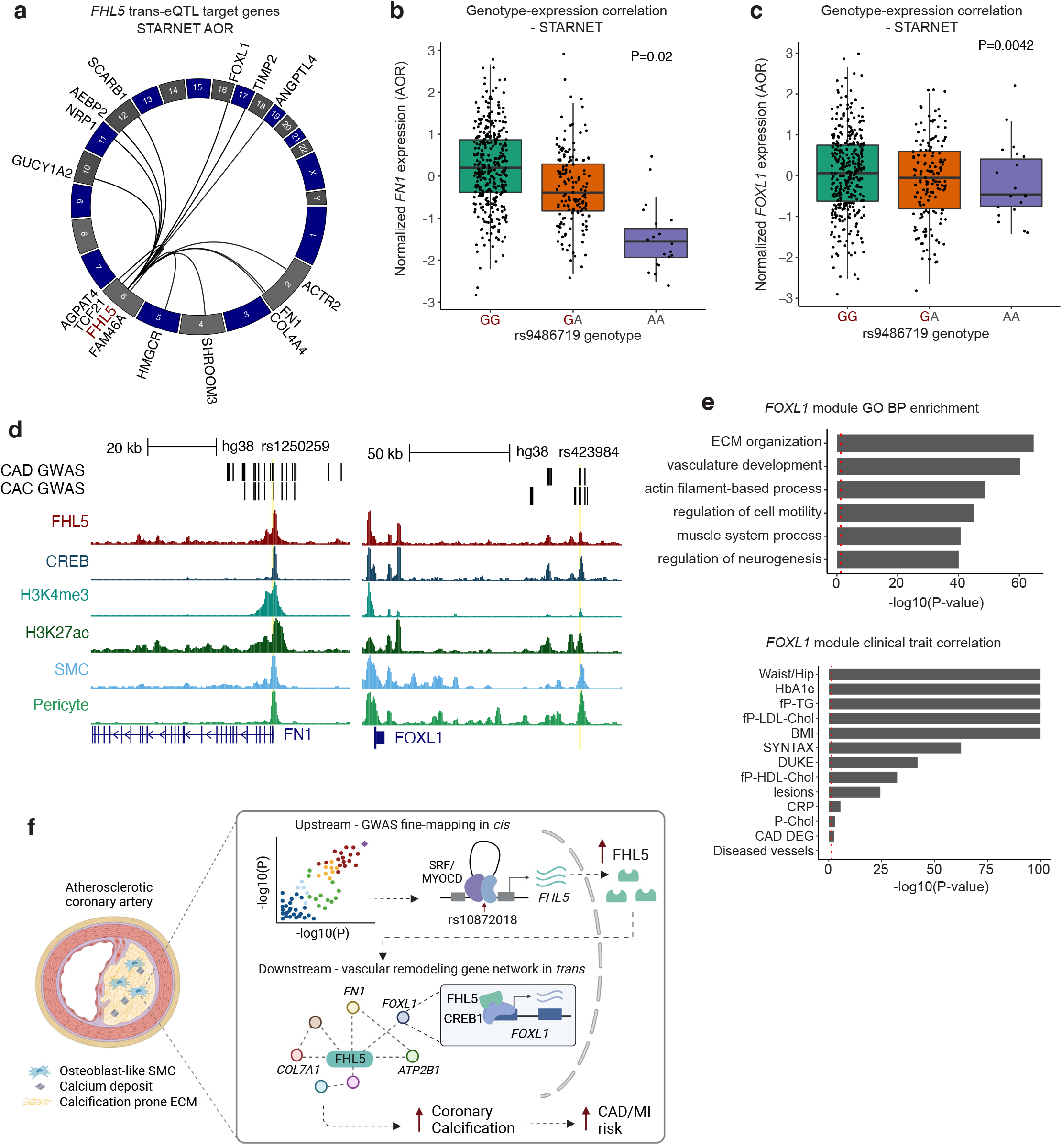
FHL5 regulates a network of CAD associated genes in human artery tissues. **(a)** Circos plot showing top *trans*-eQTL target genes for *FHL5* lead SNP rs9486719 from STARNET artery tissues, which also harbor FHL5 binding sites in SMC. Genes are plotted according to genomic position with each black link representing significant *trans*-eQTL effects (P<0.05). **(b)** Association of rs9486719 with *FOXL1* gene expression in atherosclerotic aortic artery (AOR) in STARNET. **(c)** Association of rs9486719 with *FN1* gene expression in AOR tissue. The red allele color indicates the CAD risk allele (rs9486719-G). **(d)** UCSC genome browser tracks showing FHL5 binding sites overlapping CREB, H3K4me3, and H3K27ac peaks, and accessible chromatin peaks in native SMC and pericytes (human coronary artery snATAC). The peak2Gene link indicates significant correlation of the candidate enhancer with the *FOXL1* promoter (left) and *FN1* promoter (right). **(e)** Top, enrichment of clinical traits in module 28 containing *FOXL1* and *FHL5* in STARNET cross-tissue networks. The red dotted line corresponds to P<0.05. Bottom, top GO-BP enriched in module 28. The red dotted line corresponds to a nominal threshold of FDR<0.05. These highlighted terms were subset from the full ranked list in Supp Table 13a. **(f)** Schematic depicting the proposed upstream and downstream mechanisms underlying the *FHL5* genetic association with CAD/MI and other vascular traits. Top, risk alleles for candidate causal variants (e.g. rs10872018) are associated with increased *FHL5* gene expression, which are predicted to function through SRF-MYOCD enhancers in *cis*. Bottom, this results in increased binding of FHL5 cofactor to CREB regulatory elements in vascular remodeling gene network in *trans*, ultimately leading to maladaptive ECM remodeling and vascular calcification in SMC and osteogenic-like SMC, thus increasing disease risk. Created with BioRender.com

We prioritized *FN1 and FOXL1*, two CAD/MI and CAC candidate genes, as likely downstream effectors of FHL5. The rs9486719-G CAD/MI risk allele associated with increased *FHL5* gene expression was associated with both increased *FN1* (**Figure 5B**) and *FOXL1* (**Figure 5C**) expression in *trans*. Supporting this genetic regulation, we identified an FHL5 binding site overlapping active regulatory marks harboring CAD and CAC risk variants at both the *FN1* promoter and upstream regulatory element of *FOXL1* (**Figure 5D**). Interestingly, this candidate *FOXL1* enhancer harbors rs423984, a variant in the MI 95% credible set that is also associated with CAC and *FOXL1* gene expression in muscle tissue (**Figure 5C** and **Figure S9A**). As is common for intergenic variants at GWAS loci, this variant is likely correlated with expression of both *FOXL1* and neighboring gene *FOXC2*, based on coronary artery snATAC peak2gene links (**Figure S9A)**.

To provide insights into the function of FHL5 target genes in human arteries, we queried STARNET cross tissue gene regulatory networks. *FN1* is a member of module 39 in AOR tissue, which was enriched in genes predicted to regulate ECM organization and cell adhesion and correlated with clinical traits indicative of CAD severity (DUKE and SYNTAX score) (**Table S12**). Similarly, we identified *FOXL1* and *FHL5* in module 28, a cross-tissue GRN that also includes CAD-associated genes, *SVEP1* and *MFEG8*. This module was functionally enriched in genes related to ECM organization, SMC contraction and calcium ion regulation (**Figure 5E, Figure S9B**, and **Table S13**), and correlated with atherosclerotic lesion presence and disease severity (**Figure 5E**). These enrichment analyses recapitulate the established role of *FN1* in the vessel wall and its link to disease and highlight *FOXL1* as a candidate effector gene during vascular remodeling.

To further explore the potential link between *FHL5* and *FOXL1*, we observed *FOXL1* was highly correlated with *FHL5* gene expression in human coronary arteries (**Figure S9C**). Consistent with the expression profile of *FHL5, FOXL1* gene expression is highly enriched in SMC and pericytes (**Figure S9D**) and in bulk artery tissues in both STARNET and GTEx (**Figure S9E** and **S9F**). Consistent with a role in adverse vascular remodeling, *FOXL1* gene expression was elevated in atherosclerotic aortic tissues relative to healthy controls (**Table S14**). Taken together, FHL5 mediated regulation of *FOXL1* and other ECM genes in *trans* may contribute in part to the mechanistic link between regulatory variants and vascular disease risk (**Figure 5F**).

## Discussion

Despite the identification of over 200 loci associated with CAD, the mechanisms underlying these associations are largely unknown. Given at least half of these loci are predicted to function independently of traditional risk factors^58^, characterization of these novel loci may provide insights into some of the ‘hidden’ heritable risk factors. Our study addresses this gap by implicating *FHL5* as the top candidate gene at the *UFL1-FHL5* locus associated with vascular diseases. *UFL1-FHL5* is also a well-established risk locus for migraine^59^, consistent with the genetic correlation and vascular etiology^60,61^. In this study, we elucidated the upstream regulatory mechanisms and downstream function of FHL5 in the human coronary artery. We showed that FHL5 promotes SMC calcification and contraction through dysregulation of vascular remodeling and calcium handling genes. Lastly, we characterized the FHL5 regulated SMC transcriptional network and highlighted *trans*-acting mechanisms mediated by FHL5 that contribute to the heritability of CAD and other common vascular diseases.

In atherosclerotic lesions, SMCs adopt phenotypes characteristic of other cell lineages, such as macrophages, fibroblasts, and osteoblasts^6,62^. SMCs that adopt this osteogenic phenotype participate in the deposition of intimal microcalcifications. Our findings support a mechanism where FHL5 mediated dysregulation of various metalloproteinases, collagens, and calcium signaling genes contributes to this pro-calcifying phenotype. Given the colocalization of RUNX2 and FHL5 in human coronary artery lesions, we postulate that FHL5 expression in this population of dedifferentiated SMCs alters the ECM composition to promote mineral deposition in the lesion. The significance of this calcification phenotype is underscored by the use of calcium scores in the clinic to identify patients at high risk for MI and other adverse cardiovascular events^63,64^. Despite observing increased contractility under basal conditions, which may reflect the homeostatic role of FHL5 in maintaining vascular tone, our studies under osteogenic conditions implicate FHL5 as a pro-calcifying factor as well. In support of a putative role in regulating both phenotypes, recent work by Karlöf et al^65^ reported that SMC contractile markers, which included *FHL5, MYOCD and CNN1*, were upregulated ∼4-fold in highly calcified carotid artery plaques relative to lowly calcified plaques. This upregulation of SMC markers also correlated with increased expression of recently characterized SMC calcification markers (e.g. proteoglycan 4 (*PRG4*)^66^. This is further supported by gene ontology enrichment analysis highlighting calcium signaling, cytoskeletal rearrangements, and muscle contraction as overrepresented pathways among differentially expressed genes between highly and lowly calcified lesions^65^. To reconcile these phenotypes, we speculate that FHL5 regulation of intracellular calcium ion homeostasis contributes to both processes, given the critical role of calcium in the initiation of SMC contraction^67,68^, cell stiffness^69,70^, and vascular calcification^36,71^.

This study confirms and extends previous work identifying FHL5 as a cofactor for CREB. Our CUT&RUN data supports a similar mechanism where FHL5, through interactions with CREB, regulates target genes associated with vascular remodeling in SMCs. Previous studies have linked activation of the CREB/ATF3 signaling pathway to increased SMC proliferation, migration, and calcification^72^. In addition to the CRE motif, we also observed strong enrichment of the AP1 binding motif in FHL5 binding sites. This motif was also found to be enriched in accessible regulatory elements throughout the SMC transition to a fibroblast-like state^73^. Although we cannot rule out additional direct interactions with members of the AP1 family, this enrichment may also reflect epigenetic functions, similar to AP1 interactions at downstream CAD-associated loci, *SMAD3* and *CDKN2B-AS1*^*74*^. Lastly, while we highlight that *FHL5* is regulated by SRF-MYOCD in *cis*, whether it directly interacts with or competes with MYOCD for binding to SMC gene promoters to regulate gene expression in *trans*^*75*^, requires further investigation.

Dissecting the regulatory network of disease associated transcription factors such as KLF4 and TCF21, have identified regulatory interactions with downstream CAD loci that govern SMC phenotypic modulation^5,6^. In support of the proposed omnigenic model, these studies highlight *trans*-acting mechanisms that may contribute to the majority of CAD heritability^76,77^. Our findings support the role of FHL5 as a “peripheral” transcriptional regulator of downstream “core” genes that have predicted functions in the vessel wall. By integrating STARNET vascular tissue *trans*-eQTL data, we identified a network of putative core genes that contribute to CAD pathogenesis, which include the well-characterized matrisome gene, *FN1* and the transcription factor, *FOXL1. FN1* has already been implicated as a regulator of SMC phenotypic modulation associated with CAD^6,78,79^. In contrast, the precise mechanism of how FOXL1 functions in artery tissues and atherosclerosis has not been fully characterized. Interestingly, *FOXL1* and neighboring gene *FOXC2* are both associated with CAD and bone mineral density^80^. As supported by our coronary artery SMC and STARNET cross-tissue GRNs, we propose that FOXL1 may regulate vascular remodeling pathways *in vivo* to impact disease risk. Functional dissection of other FHL5 target genes may uncover novel mechanisms and candidate genes contributing to heritable risk for CAD and other vascular diseases.

Although our study integrates multiple lines of human evidence to unravel the function of FHL5 in SMCs, we acknowledge known limitations. First, since we focus primarily on subclinical disease, this study does not address the potential roles for FHL5 in advanced disease stages. Future studies using more complex *in vitro* and human clinical samples will be needed to establish the causal link between FHL5 and indices of plaque stability and clinical outcomes. Given the lack of *Fhl5* expression in murine artery tissues, investigations utilizing traditional models of atherosclerosis and vascular calcific disease may not be applicable. Second, this study focuses primarily on the transcriptional role of FHL5 in SMCs. Other FHL family members, FHL1 and FHL3, are known to function as scaffolds to regulate sarcomere formation in myoblasts^14,81,82^. We speculate that FHL5, due to the presence of similar conserved LIM domains, may mediate similar interactions with the actin cytoskeleton^83,84^ or focal adhesions^85,86^. While our study provides insights into the FHL5 mediated gene regulatory mechanisms, future studies will be needed to clarify its role in mediating mechanotransduction or other cell signaling events.

In summary, this work reveals a molecular mechanism by which *FHL5*, the top candidate causal gene at the pleiotropic *UFL1-FHL5* locus, modulates vascular disease risk. We propose that FHL5 regulates a network of downstream genes in SMCs that participate in adverse vascular remodeling events driving disease progression. Similar to the effects of modulating other transcriptional regulators, modest increases in FHL5 gene expression may be propagated through its gene regulatory network in the vascular wall, ultimately increasing CAD/MI risk over time. Characterization of these *trans* effects in SMCs also highlights regulatory interactions at the molecular level governing atherogenic SMC phenotypic modulation. Future studies mapping genome-wide interactions of other disease associated transcriptional regulators may uncover new mechanisms and prioritize effector genes that impact primary disease processes, thereby accelerating the development of therapeutics to prevent MI and other vascular related pathologies.

## Supplementary Methods

### Coronary artery tissues and human subjects

Freshly explanted hearts from orthotopic heart transplantation recipients were procured at Stanford University under IRB approved protocols and written informed consent. Participants were not compensated for this study. Hearts were arrested in cardioplegic solution and rapidly transported from the operating room to the adjacent lab on ice. The proximal 5-6 cm of three major coronary vessels (left anterior descending (LAD), left circumflex (LCX), and right coronary artery (RCA)) were dissected from the epicardium on ice, trimmed of surrounding adipose (and in some samples the adventitia), rinsed in cold phosphate buffered saline (PBS), and rapidly snap frozen in liquid nitrogen. Coronary artery samples were also obtained at Stanford University (from Donor Network West and California Transplant Donor Network) from non-diseased donor hearts rejected by surgeons for heart transplantation and procured for research studies. All hearts were procured after written informed consent from legal next-of-kin or authorized parties for the donors. Reasons for rejected hearts include size incompatibility, comorbidities, or risks for cardiotoxicity. Hearts were arrested in cardioplegic solution and transported on ice following the same protocol as hearts used for transplant. Explanted hearts were generally classified as ischemic or non-ischemic cardiomyopathy and prior ischemic events and evidence of atherosclerosis was obtained through retrospective review of electronic health records at Stanford Hospital and Clinics. The disease status of coronary segments from donor and explanted hearts was also evaluated by gross inspection at the time of harvest (for presence of lesions), as well as histological analysis of adjacent frozen tissues embedded in Tissue-Tek O.C.T. compound (Sakura) blocks. Frozen tissues were transferred to the University of Virginia through a material transfer agreement and Institutional Review Board approved protocols. All samples were then stored at -80°C until day-of-processing.

### GWAS fine-mapping analyses

We used Summary-level Mendelian Randomization (SMR)^28^ to prioritize candidate causal genes that underlie the vascular GWAS signals at the *UFL1-FHL5* locus. SMR is a transcriptome wide association study (TWAS) method that tests whether the association of a variant with a phenotype is mediated through changes in gene expression, while accounting for the linkage disequilibrium in the study population. We performed SMR using STARNET and GTEx artery tissue eGenes for all genes +/-500kb of the FHL5 transcription start site (*FHL5, UFL1, GPR63, NDUFAF4, KLHL32, MMS22L*), using the 1000G European reference panel. We used the HEIDI (heterogeneity in dependent instruments) test to filter associations driven by linkage disequilibrium rather than pleiotropy (pHEIDI > 0.01) as previously done^29^. We considered pSMR < 0.008 (0.05/6) and pHEIDI > 0.01 as evidence of pleiotropy or causality.

In a complementary approach, we performed a Bayesian colocalization analysis using the R package, *coloc 4.0-3* ^*26*,27^ to calculate the posterior probability that the GWAS and eQTL data share a common signal. We colocalized GWAS signals with GTEx and STARNET artery tissue eQTLs using the coloc.signals function with default priors (p1=1E-4, p2=1E-4, p12=1E-5). We considered the PP4 > 0.80 as evidence of colocalization.

We used the Bayesian fine-mapping tool, PAINTOR_V3^30^ to prioritize CAD causal variants at the *UFL1-FHL5* locus. We included 1Mb around the lead CAD GWAS SNP, rs9486719 and computed Pairwise Pearson correlations for all bi-allelic SNPs based on the 1000G European reference panel. We incorporated human coronary artery ATAC-seq peaks from Turner et al^40^ as functional annotations to calculate prior probabilities for the model. Following calculation of the posterior probability, we ranked SNPs based on this metric and identified the 95% credible set.

### Human coronary artery smooth muscle cell immortalization and characterization

Primary human coronary artery SMCs (HCASMCs) were purchased from Cell Applications and immortalized by lentiviral transduction of the hTERT-IRES-hygro construct (Addgene Plasmid #85140)^45^ in 10ng/ml polybrene. Transduced cells were selected using 400μg/ml of hygromycin (Gibco) and maintained in SMC basal medium (SmBM) supplemented with SmGM-2 SingleQuots kit (PromoCell). The maintenance of similar levels of SMC marker gene expression (e.g. SM22-alpha) was confirmed via western blot. The transcriptomes of these immortalized SMCs (HCASMC-hTERT) were characterized via RNA-seq and correlated with RNA-seq data of the parental cell line^87^. A principal component analysis was performed incorporating primary HCASMCs, HCASMC-hTERT, HEPG2, K562, and human umbilical vein endothelial cell (HUVEC) RNA-seq data (**Supplementary Fig 6**). Fastq files for the primary HCASMCs were extracted from Sequence Read Archive (SRA) (SRR7064063, SRR7058289, SRR7058290). Fastq files for HEPG2, K562, and HUVECs were downloaded from ENCODE.

### Cell culture conditions

Primary HCASMCs and HCASMC-hTERT were maintained in SMC basal medium (SmBM) supplemented with SmGM-2 SingleQuots kit (PromoCell #C-22162), which includes 5% Fetal Bovine Serum (FBS), insulin, Fibroblast Growth Factor (FGF), and Epidermal Growth Factor (EGF). HEK293T and A7r5 cells were cultured in 10% FBS (Gibco) supplemented DMEM (Sigma-Aldrich). All cell cultures were maintained at 37°C and 5% CO_2_.

### Lentivirus generation and transduction

FHL5 cDNA was amplified from genomic DNA and cloned into the lentiviral vector, Lenti-III-EF1α (ABM #LV043) using EcoR1 and BamH1 restriction sites. The transfer vector, psPAX, (Addgene#12260), and pMD2.G (Addgene #12259) were transfected into HEK293T cells using Lipofectamine 3000 (Invitrogen) according to the manufacturer’s protocol. The lentiviral containing supernatant was harvested at 24 and 48 hours post transfection and frozen at -80°C until use.

HCASMCs/HCASMC-hTERT were transduced at 60% confluence with 500µl of lentiviral containing supernatant and 10ng/ml polybrene (EMD-Millipore). The viral containing supernatant was removed the following day. After 48 hours, 1.0μg/ml puromycin (Gibco) was added to the culture medium for antibiotic selection. Puromycin resistant cells were expanded, and overexpression was validated via qPCR and/or western blot.

### Immunofluorescence

Coronary arteries were embedded in OCT and cryosectioned at 8μm. Frozen slides were washed with sterile PBS twice for 2min followed by fixation with formaldehyde at 4% for 10 min. Then slides were washed with PBS twice for 2 min and tissue were permeabilized with Triton X at 0.05% for 10 min. Coronary artery tissues were blocked with donkey serum at 10% for 1 hour followed by incubation overnight at 4°C with anti-rabbit-FHL5 (Novus, NBP-32600) at 1:100 dilution and anti-mouse-α-SMA (Santa Cruz Biotechnology, SC-53142) at 1:100 dilution. Slides were washed with PBS-tween at 0.1% 3 times for 3 min each followed by incubation with secondary antibody (1:400) and F-actin (ActinGreen™ 488 ReadyProbes, ThermoFisher Scientific, USA) for 1 hr at room temperature. Then slides were washed with PBS-t (PBS, 0.05% Tween 20), 4 times for 3 min each and slides were mounted with diamond mounting medium containing DAPI. Slides were visualized with the Leica TCS SP8 confocal microscopy station and micrographs were digitized with the Leica Application Suite X software.

### Luciferase reporter assay

The enhancer sequence (180bp) harboring the rs10872018-G or rs10872018-A was cloned into the pLUC-MCS (Agilent) vector. 400ng of the luciferase reporter constructs (pLuc-MCS, pLuc-MCS-rs10872018-G, pLuc-MCS-rs10872018-A), 100ng of each expression plasmid (pCGN-SRF, pCDNA6-MYOCD, or pCMV6-empty), and 5ng of Renilla-luciferase were transfected into A7r5 cells using Lipofectamine 3000. (Invitrogen). The media was replaced 6 hours post transfection. Samples were prepared using the Dual Luciferase Assay Kit (Promega) according to the manufacturer’s protocol 24 hours post transfection. The luciferase activity was measured using the SpectraMax L luminometer (Molecular Devices). The Firefly luciferase signal was normalized to the Renilla luciferase signal for each sample.

The FHL5 bound enhancer sequence was amplified from HCASMC-hTERT genomic DNA and cloned into the lentiviral luciferase reporter plasmid, pLS-MP-Luc (Addgene #106253).

HCASMC-hTERT were transduced with 500µl of lentiviral-containing supernatant and the media was replaced the next day. Samples were prepared using the Dual-Glo Luciferase Assay system (Promega) 48 hours post transduction and the Firefly luciferase signal was normalized to total protein concentration determined using a Bradford colorimetric assay (Thermo Scientific 23225).

### Collagen gel contraction assay

250,000 HCASMCs were embedded in a 2mg/ml collagen solution (Advanced BioMatrix #5074). Following incubation at 37°C for 90 minutes, the solidified gels were detached and 1ml of media was added on top of each collagen lattice. After 18-24 hours at 37°C, the diameter of the collagen gels was measured and quantified using ImageJ.

### SMC calcification and alizarin red staining

30,000 HCASMC-hTERT were maintained in DMEM (Sigma #D6429) supplemented with 10% FBS (Gibco #26140079), 10mM β-glycerophosphate disodium salt hydrate (Sigma #G9422), 50μg/mL ascorbic acid (Sigma, #A8960), and 10nM dexamethasone (Sigma, #D2915) for 21 days, with media replaced every other day. Following 14 and 21 days of treatment, RNA was extracted from cells for qPCR. Following 21 days of treatment, cells were washed 2x with 500μL of PBS and fixed with 4% paraformaldehyde (Thermo Scientific #28908) at room temperature for 15min. Wells were rinsed twice with water and incubated with 500μL of alizarin red staining solution (Sigma, #TMS-008-C) for 20min at room temperature. Excess dye was removed by rinsing with water. Alizarin red staining was quantitated by extraction of stained calcified material at low pH. 400μL of 10% acetic acid (Ricca Chemical #R0135000) was added to each well and incubated at room temperature for 30 min. The monolayer and acetic acid slurry was vortexed, heated to 85°C for 10 min, cooled on ice for 5 min, and centrifuged at 4°C at 20,000xg for 15 min. The supernatant was neutralized with 150µl of 10% ammonium hydroxide (Alfa Aesar 35575-AP) and OD405 was measured. Alizarin red concentration was determined relative to known concentrations of Alizarin Red standards.

### SMC calcium quantification

SMCs were incubated with Fluo-4 AM (2.5 μM; ThermoFisher #F14201) and pluronic acid (0.004%) at 37°C for 20 minutes^88^. Images were then acquired at 30 frames per second using an Andor Revolution WD (with Borealis) spinning-disk confocal imaging system (Andor Technology) comprising an upright Nikon microscope with a 40X water-dipping objective (numerical aperture, 0.8) and an electron-multiplying CCD camera. Fluo-4 was excited using a 488 nm solid-state laser, and emitted fluorescence was captured using a 525/36 nm band-pass filter. Following baseline measurements, SMCs were treated with phenylephrine (5 µM; Sigma). Ca^2+^ ionophore, ionomycin (10 μM; Sigma #I0634) was used at the end of the experiment to calculate the total maximum calcium concentration^89^.

Images were analyzed using custom-designed SparkAn software^90,91^. Fractional fluorescence traces (F/F_0_) were obtained by region of interest (ROI) using polygon drawing on each cell, excluding the nucleus.

Estimates of [Ca^2+^]_i_ were made using the following equation,

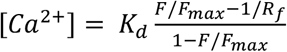

where F is fluorescence measured within an ROI, Fmax is the fluorescence intensity of Fluo-4 at a saturating maximum Ca^2+^ concentration, K_d_ is the dissociation constant of Fluo-4 (340 nM)^92^, and R*f* (= 100) is the Fluo-4 AM maximum:minimum ratio measured at saturating and zero Ca^2+^ concentrations^93^. Fmax was obtained individually for each culture dish by adding the Ca^2+^ ionophore, ionomycin (10 μM), and 20 mM external Ca^2+^ at the end of the experiment. Fractional fluorescence (F/F_0_) was determined by dividing the fluorescence intensity (F) within an ROI by the mean fluorescence value (F_0_), determined from images collected before stimulation.

### Western blot analysis

Cells were lysed in 1X RIPA buffer (Millipore Sigma #20-188) supplemented with a protease inhibitor cocktail (Roche #11697498001). Protein concentration was determined by a BCA Protein Assay (Thermo Scientific #PI23225). 20ug of protein lysate per sample was fractionated through 4-20% Tris-Glycine gel (Invitrogen #XP04205BOX) under denaturing conditions. Gels were transferred at 20V for 1hr to 0.2um pore size PVDF membrane (Invitrogen #LC2002) using the Invitrogen Mini Gel Tank (Invitrogen #A25977). The PVDF membrane was blocked for 1hr at RT in 5% non-fat dry milk (Lab Scientific #M0841) in PBS with 0.1% tween-20 (PBS-t) and incubated overnight at 4°C with primary antibody diluted in 5% non-fat dry milk. The following primary antibodies were used in western blots: anti-FHL5 (Abnova #H00009457-M01) at 1:1000 dilution, anti-HA (Abcam #ab9110) at 1:5000 dilution, anti-H3 (Cell Signaling #9715S) at 1:5000 dilution, anti-alpha smooth muscle Actin (Abcam #ab5694) at 1:2500 dilution, and anti-SM22 (Abcam #ab14106). The following day the membrane was washed 3x with PBS-T, incubated with a HRP conjugated secondary antibody (Abcam #ab205718) for 1 hour diluted 1:10000 in 5% non-fat dry milk. The signal was detected using chemiluminescent substrate (Thermo Scientific #34580) and visualized in a linear range on the Amersham Imager 600 (GE Healthcare).

### Quantitative PCR analysis

RNA was extracted using the Quick-RNA Minoprep kit (Zymo Research # R1055). Equal amounts of RNA, quantified by the Qubit Fluorometer (Invitrogen #Q33326), was reverse-transcribed into cDNA the high-capacity RNA-to-cDNA kit (Applied Biosystems #4387406) according to the manufacturer’s instructions. Taqman qPCR was performed in duplicate for each sample using the QuantStudio 5 qPCR instrument (ThermoFisher), normalized to GAPDH expression levels and analyzed via the standard 2^-ΔΔCt^ method. Alternatively, SYBR qPCR was performed in duplicate for each sample, normalized to U6 or GAPDH expression levels, and analyzed using the standard 2^-ΔΔCt^ method. All SYBR oligos were pre-validated for single amplicons using high-resolution melt curve analyses in QuantStudio.

### RNA-seq library preparation, sequencing, and analysis

#### HCASMC-hTERT

RNA was extracted from HCASMC-hTERT cells (HA, FHL5 or FHL5-NLS) using the Quick-RNA Miniprep kit (Zymo Research #R1054). The quality of RNA was assessed on the Agilent 4200 TapeStation. High quality samples, (RIN score > 8) were sequenced at the University of Virginia Genome Analysis and Technology Core in triplicate. Salmon^94^ was used to quantify transcripts from demultiplexed fastq files. Data normalization and differential expression (DE) analysis was performed using DESeq2 (v1.30.1). The Wald Test as implemented in the standard DESeq2 pipeline was used to determine differentially expressed genes. Genes with FDR < 0.05 and log2 fold change > 0.6 were considered significant. Gene ontology enrichment and pathway analyses were performed using the EnrichR web-server^95,96^ or ClusterProfiler (v4.1.4)^97,98^.

#### Coronary artery tissues

RNA was extracted from ∼50mg of frozen human coronary artery tissue using the Qiagen miRNeasy Mini RNA extraction kit (Qiagen #217004). Prior to column-based RNA isolation, the tissue was pulverized using a mortar and pestle and then homogenized in Qiazol lysis buffer using stainless steel beads in a Bullet Blender (Next Advances). RNA concentration was determined using Qubit 3.0 and RNA quality was determined using the Agilent 4200 TapeStation. Libraries were prepared from high quality RNA samples (RNA Integrity Number (RIN) > 5.5 and Illumina DV_200_ > 75) using the Illumina TruSeq Stranded Total RNA Gold kit (catalog #20020599) and barcoded with TruSeq RNA unique dual indexes (catalog # 20022371). 150bp paired end sequencing was performed at Novogene on an Illumina NovaSeq S4 Flowcell to a medial depth of 100 million total reads per library. The raw passed filter sequencing reads obtained from Novogene were demultiplexed using the bcl2fastq script. The quality of the reads was assessed using FASTQC and the adapter sequences were trimmed using trimgalore. Trimmed reads were aligned to the hg38 human reference genome using STAR v2.7.3a according to the GATK Best Practices for RNA-seq. To increase mapping efficiency and sensitivity, novel splice junctions discovered in a first alignment pass with high stringency, were used as annotation in a second pass to permit lower stringency alignment and therefore increase sensitivity. PCR duplicates were marked using Picard and WASP was used to filter reads prone to mapping bias. Total read counts and RPKM were calculated with RNA-SeQC v1.1.8 using default parameters and additional flags “-n 1000 -noDoC -strictMode” and GENCODE v30 reference annotation. The transcript and isoform expression levels were estimated using the RSEM package^99^.

### CUT&RUN assay, library preparation and sequencing

CUT&RUN assay was performed as previously described with few modifications^100^. The detailed protocol can be found at https://www.protocols.io/view/cut-amp-run-targeted-in-situ-genome-wide-profiling-14egnr4ql5dy/v3. Briefly, 250,000 HCASMC-hTERT were washed twice with wash buffer 20mM HEPES pH 7.5, 150mM NaCl, 0.5mM spermidine, 1x protease inhibitor). 10µl conA beads (Bang Laboratories #BP531) were washed three times with Binding Buffer and incubated on ice until use. Cells were incubated at RT for 10min on an orbital shaker (nutator, VWR) with the activated conA beads. The mixture was incubated overnight at 4°C with primary antibody (rabbit anti-FHL5 or anti-HA or IgG) diluted in Antibody Buffer (1x Wash Buffer, 0.05% Digitonin, 2.5mM EDTA). All primary antibodies and secondary antibodies are described in Table S2. Next day, the unbound antibody was washed away with 1mL Dig-Wash Buffer (0.05% digitonin) twice. Samples were resuspended in 150µl Dig-Wash Buffer and incubated with diluted secondary antibody for 1 hour at RT. Unbound secondary antibody was washed away with three washes of Dig-Wash Buffer and then resuspended in 50µl Dig-Wash buffer. 2.5µl of pAG-MNase (Epicypher # 15-1016) was added to each sample and incubated for 1 hour at 4°C on a nutator. After washing away unbound pAG-MNase three times with Dig-Wash buffer and resuspension in 150µl of Dig-Wash buffer, 1mM CaCl_2_ was added to activate MNase cleavage. Samples were incubated at 0°C for 30min. 150µl STOP Buffer (200mM NaCl, 20mM EDTA, 50ug/mL RNASE A, 40ug/mL glycogen) was added to halt digestion and the resulting DNA fragments was released into solution following Proteinase K digestion and purified via phenol-chloroform extraction. DNA pellets were dissolved in 30µl of TE Buffer (1mM Tris-HCl pH8.0, 0.1mM EDTA) and stored at 4°C overnight.

The CUT&RUN libraries were prepared as previously described^48,49^. 10µl of ERA buffer (4X T4 Ligase Buffer, 2mM dNTP, 1mM ATP, 10% PEG4000, 0.5U/µl T4 PNK, 0.05U/µl T4 DNA polymerase, 0.05U/µl of Taq DNA polymerase) was added to each sample. The samples were placed in a thermocycler with the following program: Cycle 1: 12°C /15min, Cycle 2: 37°C/15min, Cycle 3: 58°C/ 45min, Cycle 4: 8°C Hold. 5µl of 0.15uM of annealed Illumina Truseq adapters and 40µl of 2x Ligation Buffer (2x Rapid Ligase Buffer (Enzymatics #B101L) and 4µl T4 Ligase (Enzymatics L6030-HC-L) were added. Samples were then incubated at 20°C for 25min. Following adapter ligation, the libraries were amplified 14 cycles using NEBNext Q5 Ultra Master Mix (NEB #M0544L) and purified with 1.2X volume of Ampure Beads (Beckman CoulterCatalog #A63880). DNA was eluted into 20µl of Tris-EDTA Buffer and stored at -20°C until sequencing. DNA concentration was determined using Qubit dsDNA High Sensitivity Assay (Invitrogen #Q32851) and library size was assessed using DNA High Sensitivity Tape-Station kit (Agilent #5067-5584). Libraries were pooled and paired-end sequencing (2×42bp, 8bp index) was performed using the NextSeq 2000 instrument with the NextSeq 2000 P2 kit at the UVA Genomics Core.

### CUT&RUN analysis

CUT&RUN libraries were analyzed using the CUT&RUN Tools pipeline^50,101^. Briefly, adapter sequences were trimmed from demultiplexed reads using Trimmomatic^102^ and filtered reads were aligned to the hg38 genome with bowtie2^103^. After removal of unmapped and duplicate reads using Picard^104^, peaks were called using MACS2^105^ with a threshold qvalue < 0.01. The pooled IgG sample was used as the control in peak calling. The global settings in the pipeline were as described: https://github.com/fl-yu/CUT-RUNTools-2.0, aside from organism_build=hg38. The parameters for the individual software called by the pipeline were also left unchanged. Bigwig files for each sample were created using deeptools^106^ and visualized on the UCSC genome browser. Gene ontology analysis was performed through GREAT^54^ using the basal-plus extension to define putative target genes of FHL5 peak set, with the whole genome used as background. Known motif enrichment in FHL5, CREB, and IgG binding sites was determined in HOMER (v4.11) using the findMotifsGenome.pl command. FHL5 peaks were annotated according to the nearest protein-coding gene in CHIPseeker^51^. Differential expression analysis and FHL5 biding site data was integrated using the Binding and Expression Target Analysis (BETA) software package^53^. BETA calculates a rank sum product score to reflect the likelihood of direct transcriptional regulation by incorporating both DEGs and epigenomic profiles. We used the following thresholds for expression changes: log2 Fold Change > 0.6, and for binding: FDR < 0.05 and Q < 0.01 and signal value > 5. We considered the BETA rank sum product < 1E-3 as evidence of direct transcriptional regulation.

### GWAS SNP enrichment

We used the GREGOR software tool^56^ to determine the enrichment of GWAS risk variants in FHL5, CREB, and IgG binding sites. GWAS summary statistics were filtered to include SNPs P<5E-5. This list of suggestive SNPs was pruned using PLINK (v1.9) to retain the most significant SNPs with pairwise LD (r2) threshold < 0.2 in the 1000G European reference panel. We then used the default GREGOR parameters as described in https://genome.sph.umich.edu/wiki/GREGOR.

### STARNET analyses

#### eQTL analysis

The STARNET cohort and datasets were described previously^25,107^. The STARNET cohort includes 600 individuals with CAD and 250 control samples. RNA-seq libraries from cardiometabolic tissues (aorta, mammary artery, liver, subcutaneous fat, visceral fat, blood, and skeletal muscle) were prepared using the polyA and Ribo-Zero library preparation protocols. 50-100bp single-end sequencing was performed using the Illumina HiSeq sequencer to a depth of 20-30million reads per sample. *Cis*-eQTLs and *trans*-eQTLs were identified using the Matrix QTL package.

#### Gene regulatory network analysis

Normalized gene expression for all STARNET tissues were used to construct co-expression modules using block-wise Weighted Gene Co-expression Network Analysis (WGCNA). The modules were annotated based on gene ontology enrichment analyses using the Fisher’s exact test employed in the WGCNA package. Phenotypic correlations with clinical traits were calculated by aggregating Pearson correlation *P-*values for genes in the module by Fisher’s method. Gene regulatory networks were inferred using the GENIE3 package with edges constrained by eQTL genes and transcription factor annotations. Key driver analyses were performed in the Mergenomics R package. The co-expression and gene regulatory network data can be accessed at http://starnet.mssm.edu/.

### Coronary artery gene regulatory network analysis

#### WGCNA

Total RNA-seq data was filtered to exclude genes not present in greater than 80% of the samples. Hierarchical clustering was used to identify sample outliers (UVA047, UVA125). The remaining 15,720 genes and 148 samples were used as input into WGCNA^41^ to detect gene modules. WGCNA calculates the co-expression of genes through an adjacency matrix based on co-expression similarity between the i-th gene and the j-th gene. Hierarchical clustering of these gene co-expression values was used to determine gene modules. Module detection was performed multiple times using iterativeWGCNA^42^ to prune poorly fitting genes and generate more robust gene modules. Genes not assigned to any of the modules were designated to the gray module. Coexpression modules were visualized using Cytoscape.

#### Bayesian network construction

Bayesian networks for the *FHL5*-containing module were constructed using RIMBANET (Reconstructing Integrative Molecular Bayesian Networks)^108^. STARNET aortic tissue eQTL data were also used as genetic priors such that genes with *cis*-eQTLs are allowed to be parent nodes of genes with coincident trans eQTLs, but not vice versa. One thousand Bayesian networks were reconstructed using different starting random seeds. Edges that appeared in greater than 30% of the networks were used to define a consensus network.

#### Key driver analysis

Given a set of genes (G) and directed gene network (N), Key driver analysis (KDA) generates a subnetwork, N_G, defined as the set of nodes in N that are no more than h-layers from the nodes of G and subsequently computes the size of the h-layer neighborhood (HLN) for each node. The key driver score increases if the HLN was greater than μ + *σ*(μ), where μ is average size of HLN. Total key driver scores for each node were computed by summing all scores at each h-layer scaled according to h, 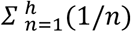. Genes with key driver scores in the top 5% were annotated as module key drivers.

### Statistical analyses

Data in bar graphs are presented with mean+/- standard error of mean (SEM) with each point represented as an individual replicate. Data in box plots are presented with lines denoting the 25th, median and 75th percentile with each point representing an individual donor. Pairwise comparisons were made using the student’s t test or Wilcoxon rank test as appropriate. Comparisons between more than two groups were assessed using a one-way ANOVA test or Kruskal-Wallis test. The normality of the data was assessed using the Shapiro-Wilkes test, with P > 0.05, supporting a normal distribution. For each of these analyses, we considered P < 0.05 as significant.

To identify differentially expressed genes, we used the false discovery rate (FDR) adjusted P< 0.05 threshold and log2FoldChange > 0.6. Heatmaps were created using the pheatmap package and represent normalized expression (Z-score) for genes, scaled across each row. Gene ontology enrichment analyses were performed relative to all expressed genes using Fisher’s Exact Test, with a significant threshold of 5% FDR.

## Acknowledgements

We thank Dr. Pat Pramoonjago and Sheri VanHoose and members of the histology core facilities for tissue sectioning and histological staining assistance and Dr. Katia Sol-Church and members of the Genomics Core facility for assistance with library construction and sequencing. We also thank Dr. R. Kirk Riemer and Dr. Xiaoyuan Ma at Stanford University for helpful discussions.

## Sources of Funding

This work was supported by grants from: the National Institutes of Health (R01HL148239 and R00HL125912 to C.L.M.; F31 HL156463 to D.W.) and the Fondation Leducq (‘PlaqOmics’ 18CVD02 to J.LM.B. and C.L.M.)

## Author Contributions

C.L.M supervised research primarily related to the study. R.M., J.LM.B., J.C.K., M.C., S.K.S. and A.M. supervised research secondarily related to the study. C.L.M. and D.W. conceived and designed the experiments. D.W., G.A., C.L.L.C, A.W.T., Y.C., M.Ku., C.J.D. and M.P. performed the experiments. D.W. and C.Y. performed the statistical analyses. D.W., G.A., Li.M., R.N.P., R.A., J.V.M., M.Ku. and M.D.K. analyzed the data. M.Ka., P.P., Lj.M., U.H., J.C.K. and J.LM.B. contributed reagents/materials/analysis tools. D.W., G.A. and C.L.M. wrote the paper.

## Disclosures

Johan Björkegren is a shareholder in Clinical Gene Network AB and has an invested interest in STARNET. Jason Kovacic is the recipient of an Agilent Thought Leader Award (January 2022), which includes funding for research that is unrelated to the current manuscript. All other authors declare that they have no competing interests relevant to the contents of this paper to disclose.

## Data Availability

All raw and processed CUT&RUN and RNA-seq datasets are made available on the Gene Expression Omnibus (GEO) database (accession: GSE201572).The following publicly available datasets were used in this study: human coronary artery scATAC-seq generated by Turner et al. (accession: GSE175621), bulk human coronary ATAC-seq generated by Miller et al. (accession: GSE72696), bulk left ventricle and liver tissue ATAC-seq generated by the ENCODE project (https://www.encodeproject.org/). Bulk RNA-seq fastq files for human coronary artery SMCs (SRR705828, SRR705829, SRR706406) were downloaded from sequence read archive (SRA) using sratools. ENCODE bulk RNA-seq fastq files for HUVECs (ENCFF000DUK, ENCFF000DUH), HEPG2 (ENCFF982FAM, ENCFF564BSM) and K562 (ENCFF104ZSG, ENCFF695XOC) cells were downloaded from the ENCODE project (https://www.encodeproject.org/). Processed scRNA-seq data for human coronary arteries generated by Wirka et al. and carotid arteries generated by Alsaigh et al. are accessible at the PlaqView single-cell data portal (https://www.plaqview.com). Normalized gene expression levels and expression quantitative trait loci (eQTL) data are available at the Genotype-Tissue Expression (GTEx) portal website (https://www.gtexportal.org). Stockholm-Tartu Atherosclerosis Reverse Network Engineering Task (STARNET) eQTL data and GRNs are available at http://starnet.mssm.edu. Summary statistics and gene annotations for cardiometabolic GWAS (hypertension, diastolic blood pressure, CAD, systolic blood pressure, BMI, and pulse pressure) were accessed through the Cardiovascular Disease Knowledge Portal (https://cvd.hugeamp.org/).

## Code Availability

All custom scripts used are available at https://github.com/MillerLab-CPHG/FHL5_Manuscript. All software tools used in this study are publicly available and full names and versions are provided in the reporting summary.

## Notes

### Competing Interest Statement

Johan Bjorkegren is a shareholder in Clinical Gene Network AB and has an invested interest in STARNET. Jason Kovacic is the recipient of an Agilent Thought Leader Award (January 2022), which includes funding for research that is unrelated to the current manuscript. All other authors declare that they have no competing interests relevant to the contents of this paper to disclose.

