## Supplemental Figures for "FHL5 controls vascular disease-associated gene programs in smooth muscle cells"

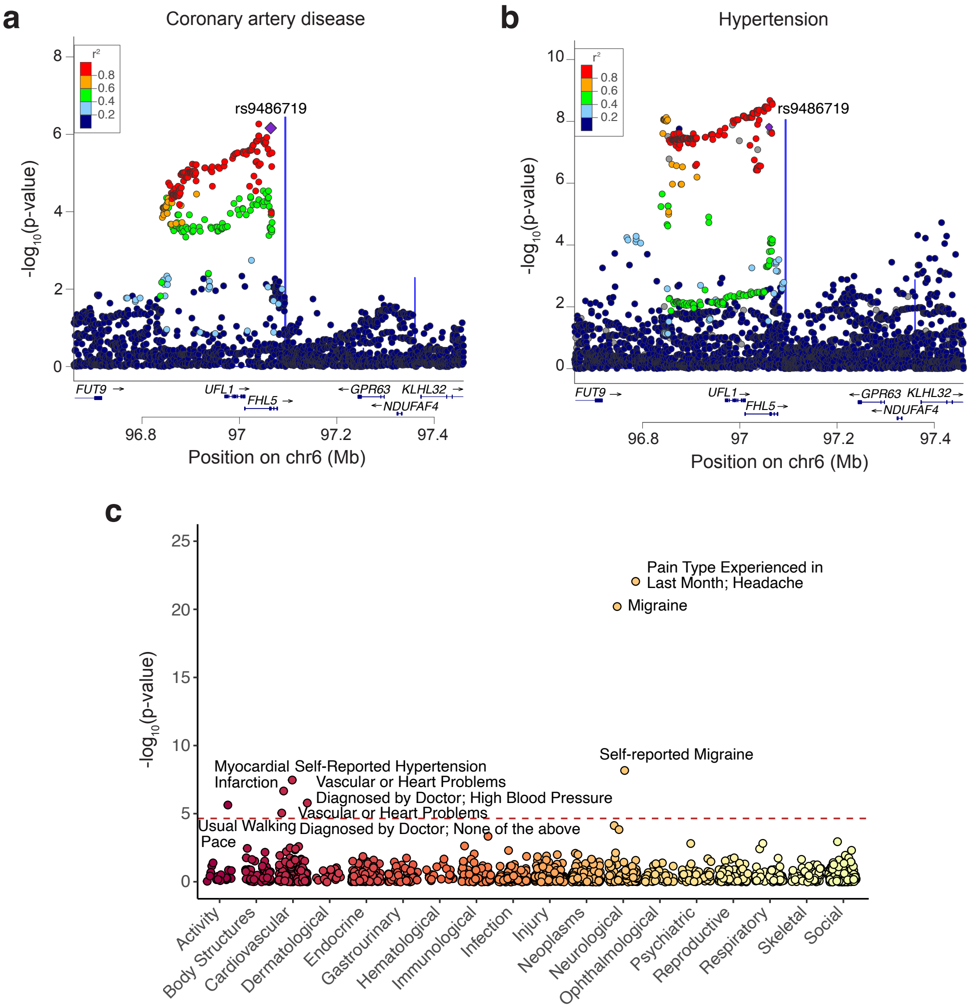


**Fig. S1. *UFL1-FHL5* locus is associated with vascular diseases.** LocusZoom plot highlighting the genetic association of the *UFL1-FHL5* locus with coronary artery disease (CAD) (a) as determined from the CARDIoGRAMplusC4D and UK Biobank meta-analysis, and hypertension (b) as detemined from the UK Biobank. (c) Genetic association of rs9486719 across multiple phenotypes using GWAS studies reported in Phenoscanner, and traits organized in physiological categories on x-axis.

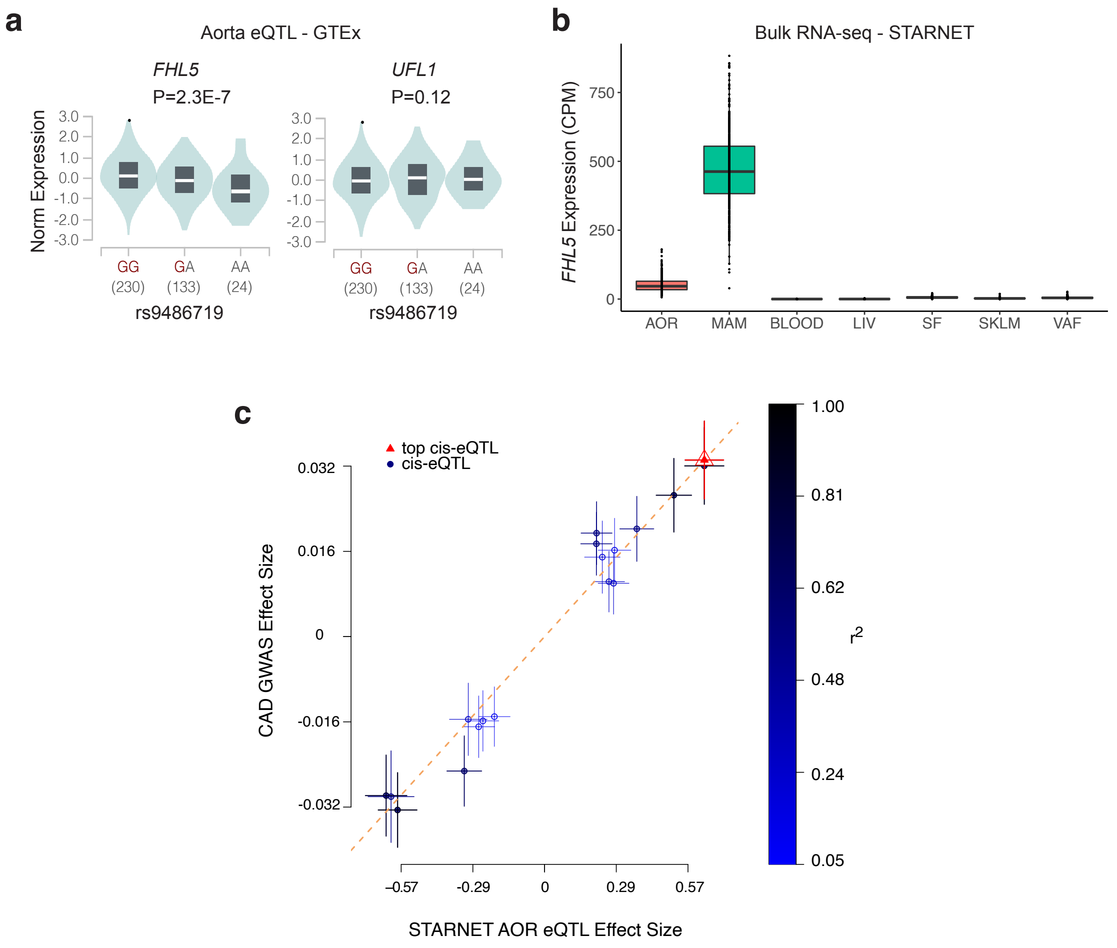


**Fig. S2. *FHL5* is the top candidate causal gene at the CAD/MI *ULF1-FHL5* locus.** (a) Violin plots showing the association of rs9486719 with *FHL5* gene expression (left) and lack of association with *UFL1* gene expression (right) in GTEx (Aorta). (b) Normalized *FHL5* gene expression in counts per million (CPM) across STARNET cardiometabolic tissues: AOR: atherosclerotic aorta, MAM: mammary artery, LIV: liver, SF: subcutaneous fat, SKLM: skeletal muscle: VAF: visceral adipose fat. (c) Correlation of the effect size of MI risk alleles with *FHL5* eQTL effect size in STARNET AOR. Color scale depicts Pearson's r-squared values for the correlation of individual alleles.

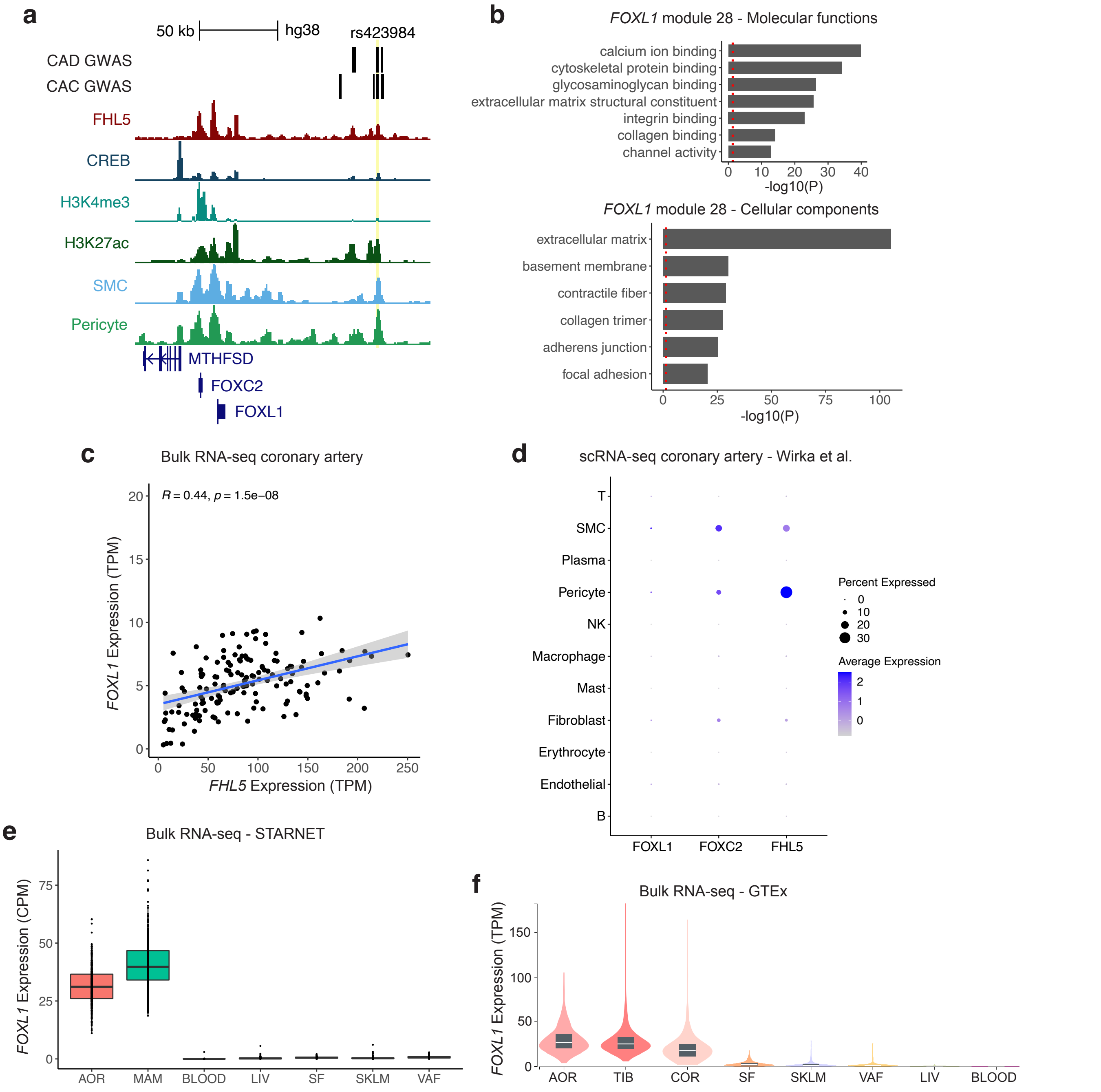


**Fig. S9. FHL5 regulation of *FOXL1* may contribute to the underlying mechanism of its association with CAD/MI.** (a) UCSC genome browser screenshot of the *FOXC2-FOXL1* locus, highlighting an FHL5 binding site upstream of the *FOXL1* promoter harboring CAD and CAC SNPs. Top credible SNP rs423984 (95% credible set for MI) highlighted in yellow. (b, upper) Top molecular functions enriched in module 28. (b, lower) Top cell compartments enriched in module 28 genes. P-values shown were adjusted using Bonferonni test for multiple comparisons. The red dotted line corresponds to a nominal threshold of FDR<0.05. (c) Pearson correlation of *FHL5* and *FOXL1* gene expression from bulk RNA-seq of human coronary arteries (N=148). (d) Dot plot showing enrichment of *FOXL1*, *FOXC2* and *FHL5* gene expression in SMC and pericytes in human coronary artery single-cell RNA-seq (Wirka et al.). (e) Normalized expression level in counts per million (CPM) of *FOXL1* in cardiometabolic tissues profiled in STARNET bulk RNA-seq analysis. (f) Normalized expression level in transcripts per million (TPM) of *FOXL1* in bulk RNA-seq of GTEx cardiometabolic tissues. AOR: aorta, MAM: mammary artery, BLOOD: whole blood, LIV: liver, SF: subcutaneuous adipose fat, SKLM: skeletal muscle, VAF: visceral adipose fat, TIB: tibial artery, COR: coronary artery.

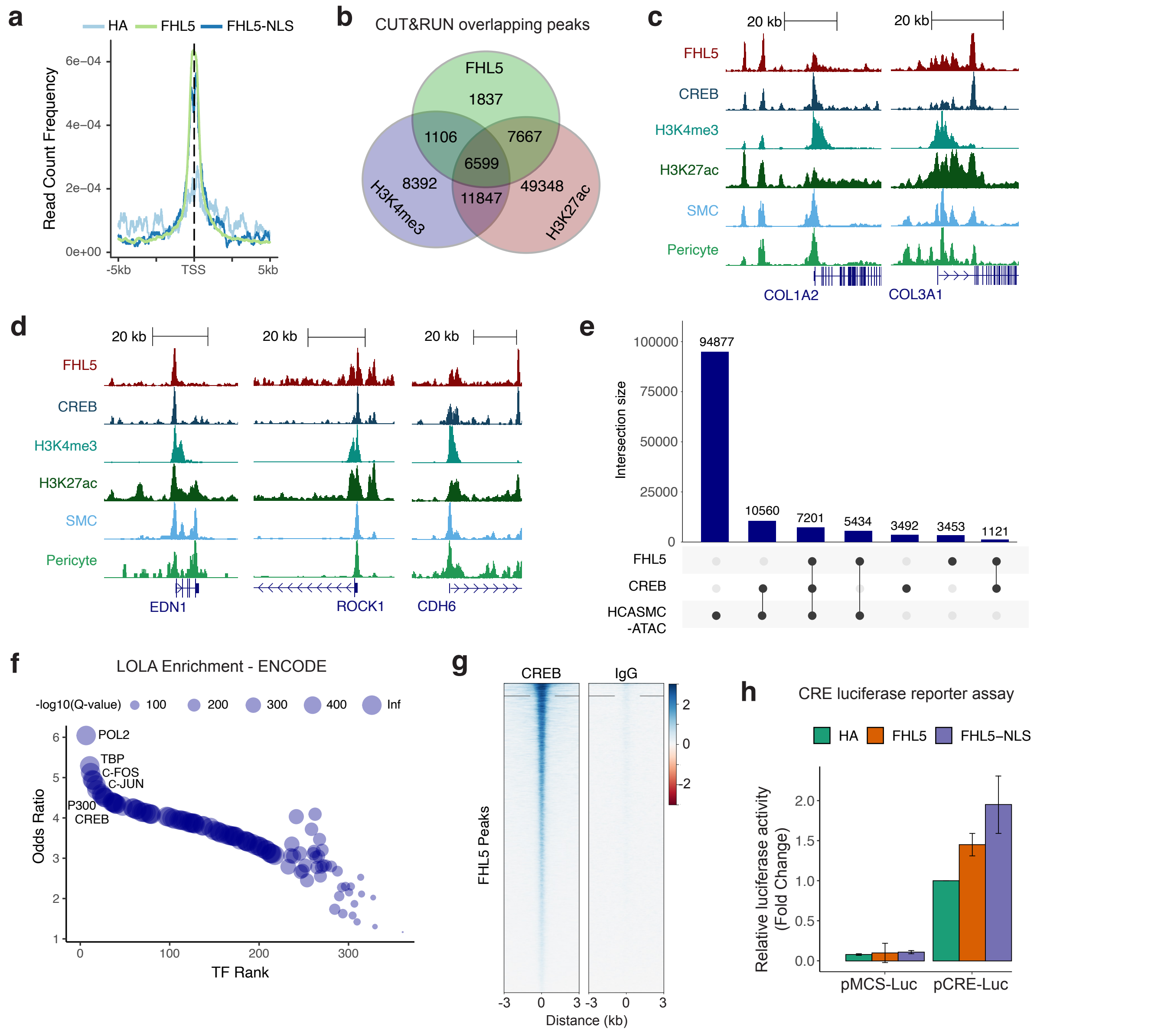


**Fig. S8. FHL5 interacts with CREB to transcriptionally regulate extracellular matrix and cell adhesion genes.** (a) Density plot showing similar enrichment of FHL5 (light green) and FHL5-NLS (dark blue) CUT&RUN peaks centered on transcription start sites (TSS). (b) Overlap of FHL5, H3K27ac, and H3K4me3 CUT&RUN peaks in SMCs. (c) UCSC genome browser tracks showing FHL5 and CREB binding sites near extracellular matrix genes, *COL1A2*, and *COL3A1*. (d) UCSC genome browser tracks showing FHL5 and CREB binding sites near the promoters of cell adhesion genes, *EDN1*, *ROCK1*, and *CDH6*. (e) Overlap of FHL5 and CREB SMC binding sites with primary human coronary artery SMC ATAC-seq peaks. (f) LOLA enrichment of FHL5 binding sites in ENCODE transcription factor (TF) ChIP-seq datasets, highlighting AP-1 family TFs, CREB, as well as transcription machinery proteins RNA polymerase II (POL2) and TATA binding protein (TBP), and transcriptional activator P300. (g) Heatmap visualization of distance of CREB binding sites to the center of FHL5 binding sites, compared to non-specific IgG binding sites. (h) Relative fold change in CRE-luciferase reporter activity after overexpression of FHL5 in HEK293T cells.

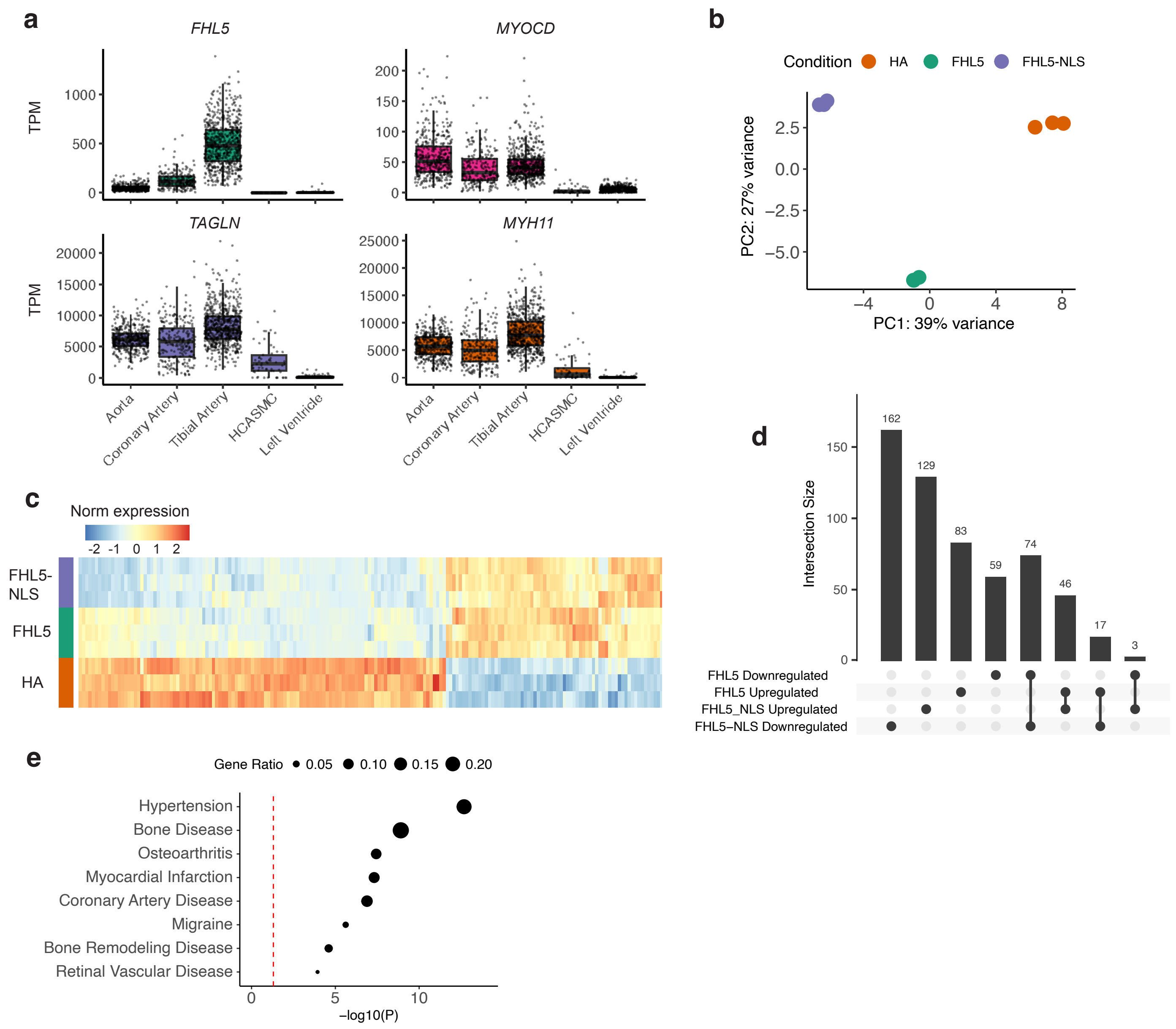


**Fig. S7. Comparison of FHL5 and FHL5-NLS differentially expressed genes.** (a) Normalized expression level in transcripts per million (TPM) of *FHL5* and SMC markers in human aorta (N=432), coronary artery (N=240), tibial artery (N=663), left ventricle (N=432) and HCASMC (N=61). (b) PCA of normalized HA, FHL5, and FHL5-NLS using RNAseq transcriptomes (n=3 per group). (c) Heatmap showing log2 normalized expression of 168 DEGs derived from the union of FHL5 and FHL5-NLS significantly upregulated and downregulated DEGs and clustered using n=3 biological replicates per group. (d) Upset plot showing overlap between FHL5 and FHL5-NLS upregulated and downregulated genes. (e) Disease enrichment analysis of the FHL5 DEGs showing enrichment of disease ontology (DO) terms from 381 FHL5 DEGs against the whole transcriptome as a background. P-values shown are unadjusted.

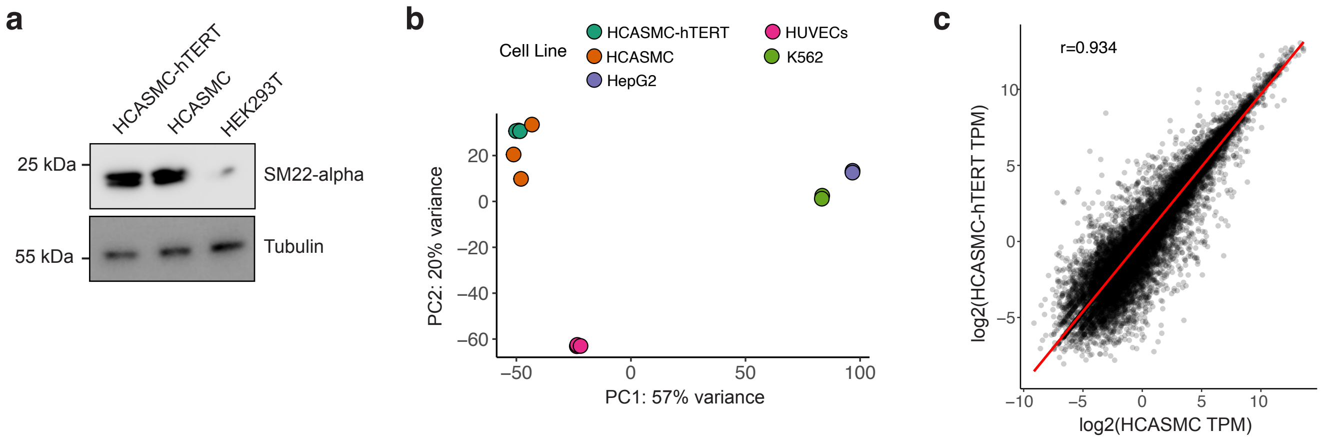


**Fig. S6. Characterization of HCASMC-hTERT.** (a) Western blot showing expression levels of SMC marker, SM22-alpha, in HCASMC-hTERT and the parental HCASMC (Cell Applications #2105). Tubulin is shown as a loading control. Molecular weights are shown for protein markers in PageRuler Plus ladder. (b) Principal component analysis (PCA) showing similarity in the transcriptomes of HCASMC-hTERT and primary HCASMC relative to HEPG2, HUVEC, and K562 cells. (c) Pearson correlation of HCASMC-hTERT and parental primary HCASMC transcriptomes shown as log2 normalized gene expression (transcripts per million (TPM).

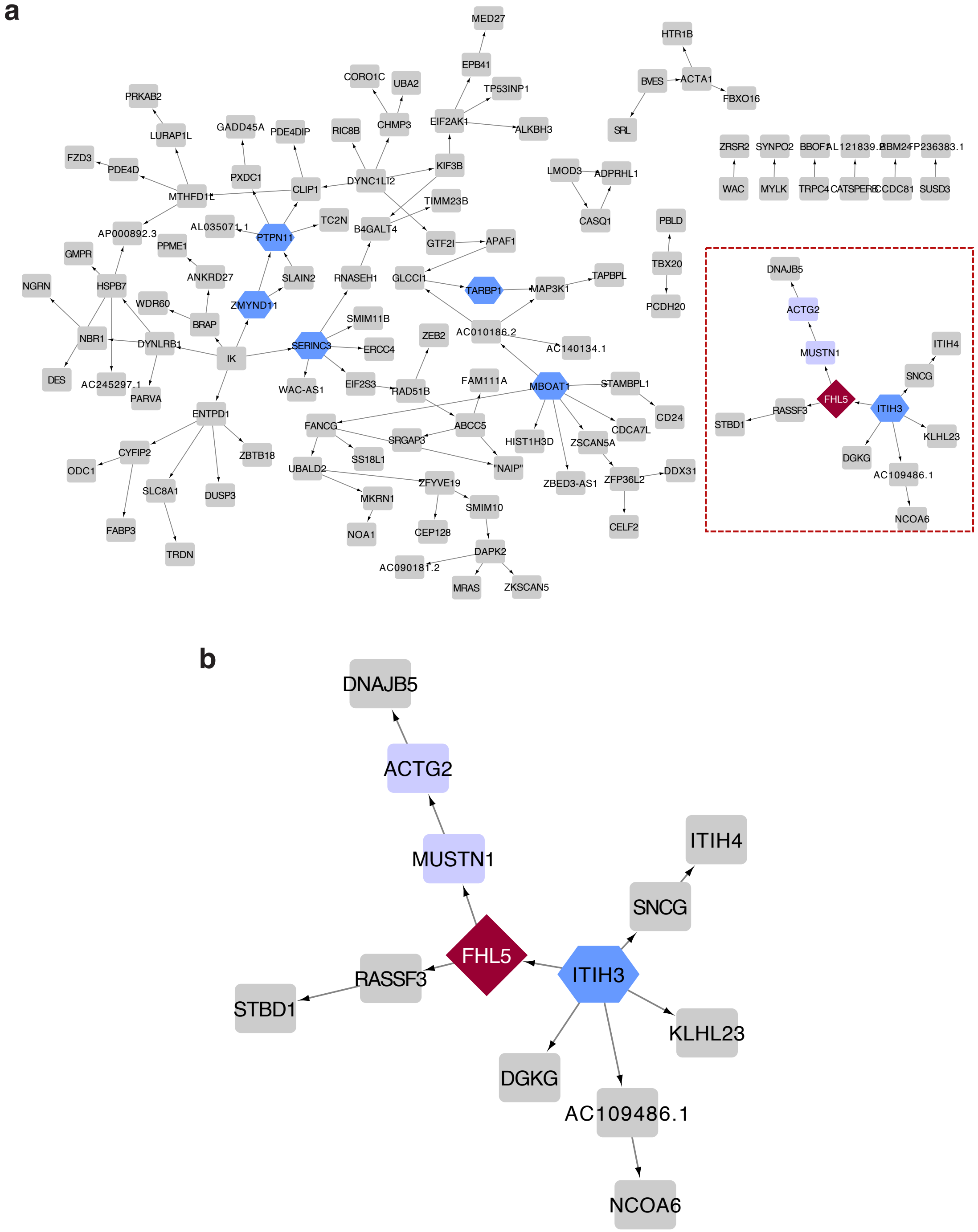


**Fig. S5.** ***FHL5*** **gene regulatory network (GRN) in human coronary arteries.** (a) Directed network depicting the *FHL5* GRN in human coronary arteries. The key drivers in the network are highlighted in blue hexagons. The dotted red box denotes the *FHL5* subnetwork. (b) Zoomed in view of the *FHL5* subnetwork. The key driver of this subnetwork, *ITIH3* is depicted in the blue hexagon. Downstream FHL5 genes that function in SMC contraction biological process are highlighted in the purple rectangles.

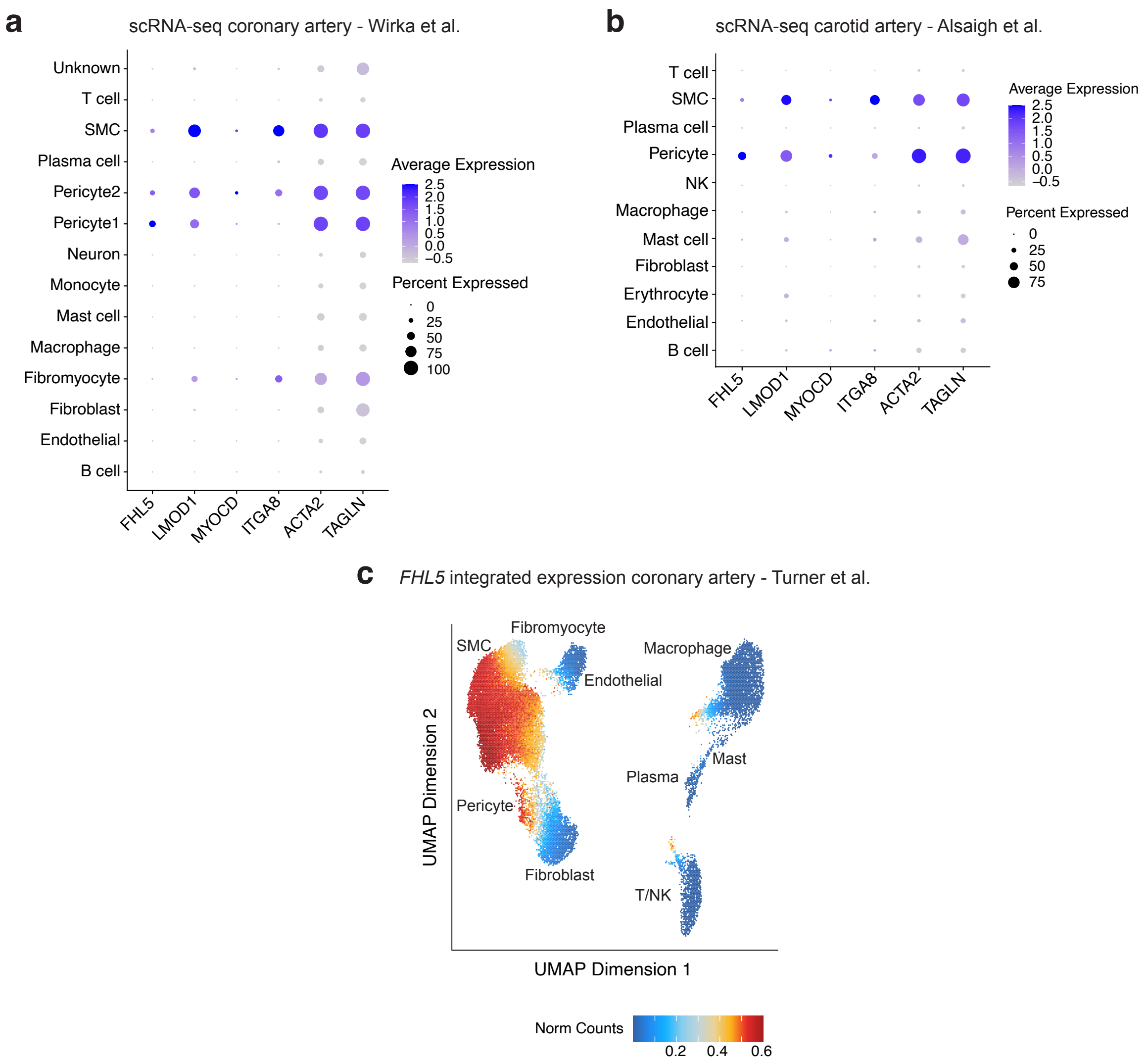


**Fig. S4. *FHL5* gene expression is enriched in SMC and pericytes in human coronary arteries.** (a) Dot plot showing average expression and percentage of cells expressing *FHL5* and other SMC markers in scRNAseq of subclinical atherosclerotic human coronary arteries (Wirka et al.). (b) Dot plot showing average expression and percentage of cells expressing FHL5 and other markers in scRNAseq of healthy and atherosclerotic human carotid arteries (Alsaigh et al.). (c) UMAP plot showing *FHL5* integrated expression in SMC and pericytes using integrated coronary artery snATAC-seq and scRNA-seq dataset from Turner et al. Clusters are labeled by the main cell-type annotations after integration of the two datasets.

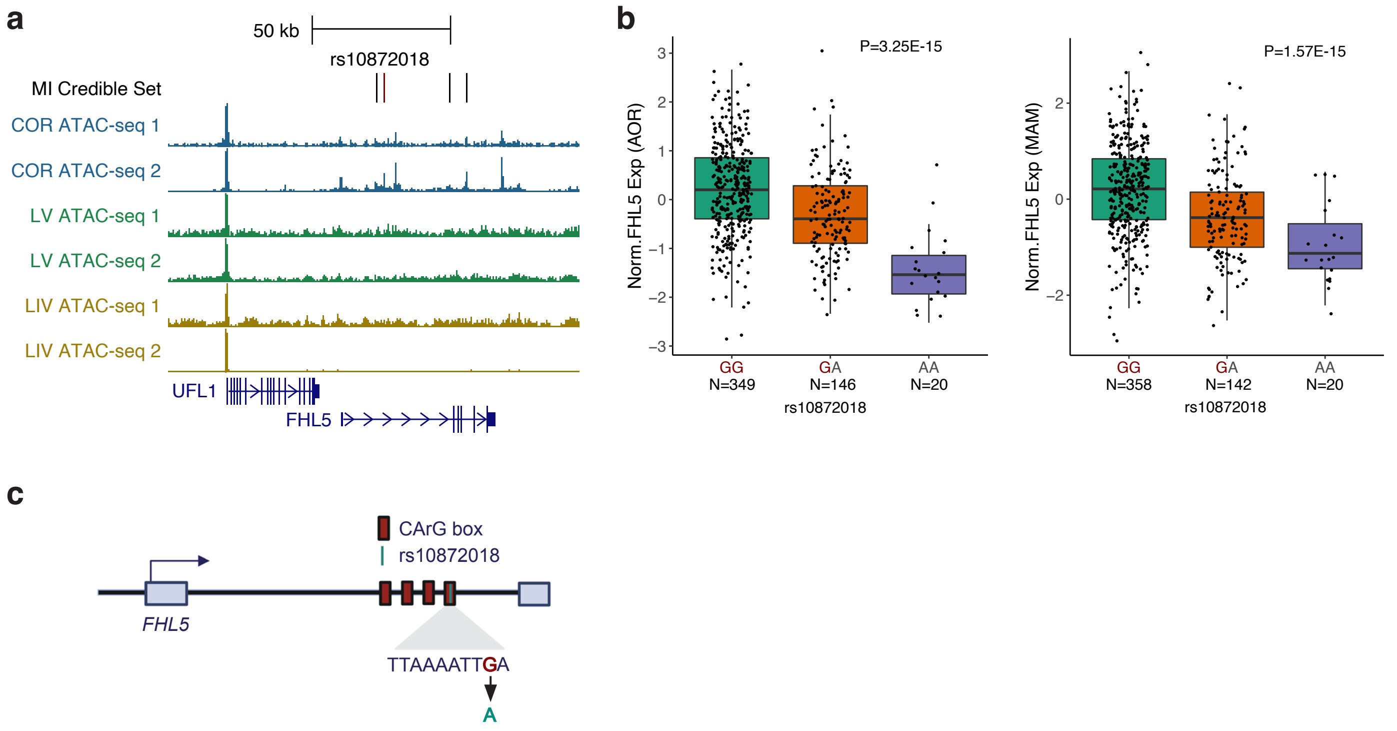


**Fig. S3. rs10872018 is the top candidate causal variant underlying the MI *UFL1-FHL5* locus.** (a) UCSC genome browser screenshot showing the overlap of *FHL5* MI GWAS credible set of variants with ATAC-seq peaks in coronary artery (COR), left ventricle (LV), and liver (LIV), from two independent donors per tissue. Top candidate SNP rs10872018 highlighted in red. (b) Association of rs10872018 with *FHL5* gene expression in STARNET atherosclerotic aorta (AOR) (left) and mammary artery (MAM) (right). (c) Schematic highlighting the 4 CArG boxes in intron 1 of *FHL5*, with the last motif disrupted by rs10872018-A.
